# Estimating the *cis*-heritability of gene expression using single cell expression profiles controls false positive rate of eGene detection

**DOI:** 10.1101/2025.02.24.639892

**Authors:** Ziqi Xu, Arya Massarat, Laurie Rumker, Melissa Gymrek, Soumya Raychaudhuri, Wei Zhou, Tiffany Amariuta

## Abstract

For gene expression traits, *cis*-genetic heritability can quantify the strength of genetic regulation in particular cell types, elucidating the cell-type-specificity of disease variants and genes. To estimate gene expression heritability, standard models require a single gene expression value per individual, forcing data from single cell RNA-sequencing (scRNA-seq) experiments to be “pseudobulked”. Here, we show that applying standard heritability models to pseudobulk data overestimates gene expression heritability and produces inflated false positive rates for detecting *cis*-heritable genes. Therefore, we introduce a new method called scGeneHE (<u>s</u>ingle <u>c</u>ell <u>Gene</u> expression <u>H</u>eritability <u>E</u>stimation), a Poisson mixed-effects model that quantifies the *cis*-genetic component of gene expression using individual cellular profiles. In simulations, scGeneHE has a consistently well-calibrated false positive rate for eGene detection and unbiasedly estimates *cis*-heritability at many parameter settings. We applied scGeneHE to scRNA-seq data from 969 individuals, 11 immune cell types, and 822,552 cells from the OneK1K cohort to infer cell-type-specificity of genetic regulation at risk genes for immune-mediated diseases and trace the fluctuation of *cis*-heritability across cellular populations of varying resolution. In summary, we developed a new statistical method that resolves the analytical challenge of estimating gene expression *cis*-heritability from native scRNA-seq data.

## Introduction

Heritability, the proportion of phenotypic variance explained by genetic variation, is critical to understanding the impact of genetic effects on diseases and complex traits at a population scale^1,2,3,4,5,6^. When applied to gene expression levels, heritability measures the magnitude of genetic control on a gene’s mRNA expression in a given cell type^7,8,9,10,11^. Detecting heritable genes is key to understanding gene function: genes that are heritable and thus under genetic control can be experimentally perturbed with sequence variation to reveal a gene’s effector function in other cell types or on other genes, pathways, and even disease endotypes. Heritable genes can also be used in genetic risk models as biomarkers to help predict clinical phenotypes and an individual’s disease risk^9,12,13,14^.

Associations between genetic variants and gene expression levels via expression quantitative trait loci (eQTL) analysis provide complementary information to heritability estimation^15,16,17,18^ (**Figure 1A**). On one hand, heritability estimates the aggregate additive effect of *cis*-genetic variation on gene expression, while eQTL analysis identifies which variants are marginally associated with changes in expression levels. However, these associations are often numerous and dependent due to linkage disequilibrium^19^, which inflates the perceived genetic effect on expression. Moreover, conventional *cis*-eQTL analysis tends to identify similar variant-to-gene links across cell types^18^, while *cis*-heritability estimation can reveal quantitative differences in the magnitude of genetic regulation, even from shared causal variants. In this study, we sought to use heritability estimation as a reliable approach to clarify the cell-type-specific nature of genetic regulation of gene expression levels, especially for genes that have been previously associated with immune-mediated polygenic diseases.

**Figure 1.**
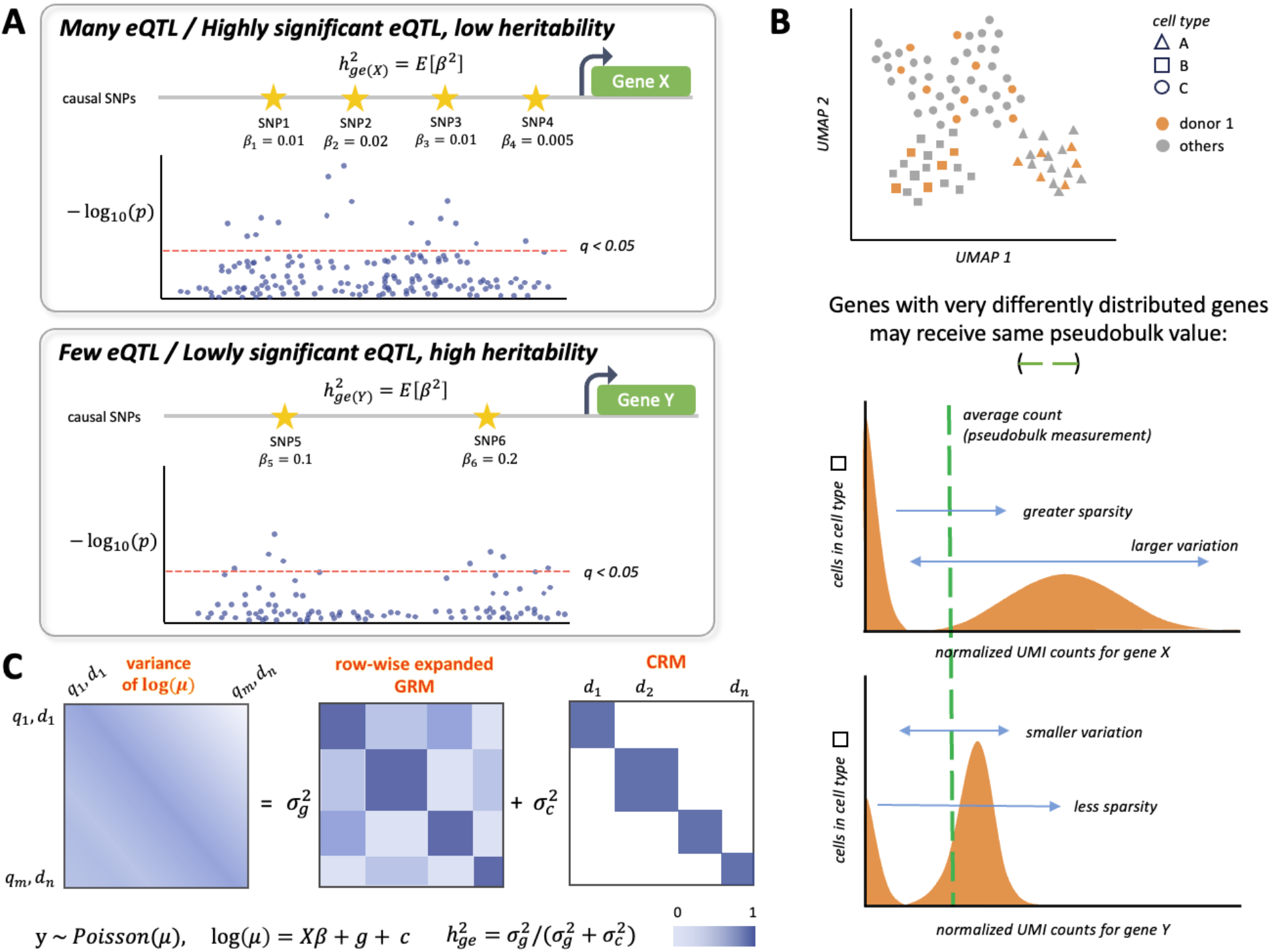
scGeneHE schematic. (**a**) *cis*-eQTL analysis and *cis*-heritability estimation provide two orthogonal pieces of information regarding the genetic architecture of gene regulation. The number of significant *cis*-eQTLs or degree of significance of those *cis*-eQTLs do not necessarily correspond to the magnitude of *cis*-heritability. *β_j_* represents the true eQTL effect size of variant *j* and *h*^2^_*ge*_ (*X*) is the *cis*-heritability of gene *X*. (**b**) Pseudobulk preprocessing loses critical information regarding the distribution of unique molecular identifier (UMI) counts for a given gene. scGeneHE leverages knowledge of this distribution to improve the certainty of *cis*-heritability estimation. (**c**) scGeneHE decomposes the log of the variance of the gene expression phenotype (a vector of all cell-donor pairs) into a shared genetic component modeled by a random effect quantifying inter-individual variation (g) and a shared cellular component modeled by a random effect quantifying intra-individual variation or cell-cell relatedness (c). Variables: q indexes cells up to m cells, d indexes donors up to n donors, 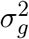 is the phenotypic variance attributed to the *cis*-genetic component of gene expression, 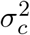 is the phenotypic variance attributed to the shared environment of the cells from a given individual, y is the Poisson-distributed gene expression profile across cells and donors, is the rate parameter for the Poisson distribution of single cell read counts, *X* are fixed effect covariates and learned coefficients, respectively. GRM, genetic relationship matrix; CRM, cell-cell relatedness matrix.

The role that genetic variation plays in regulating gene expression is highly cell-type-specific^20,21,22,23,24,25^. While cell-type-specific patterns of gene expression are now characterized at unprecedented resolution with single-cell RNA sequencing (scRNA-seq) technology, these datasets initially prioritized large numbers of cells but not donors, which is not sufficient for inferring genetic regulatory mechanisms relevant to cellular or disease-critical processes, as this relies on having substantial inter-individual genetic variation^22,26,27,28,29,30,31^. More recently, datasets have increased in donor size^27,28,32,33^. Integrative analysis of scRNA-seq with genome-wide association studies (GWAS) have proposed disease-critical cell types and cell states without explicitly modeling the genetic effects on gene expression, but rather via position-based SNP-gene linking strategies and differential gene expression analysis across cell types^34,35,25,36^. Most downstream data analysis is only compatible with bulk RNA-seq measurements, in which each donor has exactly one gene expression measurement in a given cell type for a given gene^37,38,39,40^. While scRNA-seq generates mRNA counts across a multitude of cells from each of many donors, there is no straightforward way to apply previous methods while simultaneously leveraging individual cellular expression profiles to estimate gene expression heritability: these data naturally violate the assumptions of independence between gene expression measurements which governs bulk transcriptomic analysis. In attempts to mimic the architecture of bulk RNA-seq studies, previous studies performed eQTL mapping or heritability estimation with “pseudobulk” counts generated by summing or averaging the measurements across cells of the same donor^22,24,27,28,41,42,43,44,45,46,47^. However, pseudobulk counts obscure the underlying distribution of read counts, which can provide critical information regarding a gene’s regulation (**Figure 1B**).

Other studies have creatively designed new approaches for *cis*-eQTL mapping that can explicitly model native scRNA-seq counts across cells, which recognize that normal distributions used for continuous bulk gene expression data are underpowered when applied to discrete count distributions from scRNA-seq^22,27,29,33,42,48,49,50^. Therefore, these methods employ discrete count distributions such as Poisson or negative binomial models. This work was motivated by a hypothesis that modeling intra-individual (relatedness of cells from the same individual) variation would provide greater certainty of eQTL effects and thus decrease standard errors on heritability estimation and eQTL mapping alike, relative to pseudobulk analysis. However, we found that the widely used approach of estimating heritability from pseudobulk gene expression data with GCTA^3,45^ produces frequent overestimation of *cis*-heritability and smaller than expected standard errors, resulting in high type I error for estimating non-zero *cis*-heritability (discussed in detail below). Many follow-up genomic analyses use *cis*-heritability estimates of gene expression as prior knowledge – such as gene selection and gene model building in transcriptome-wide association studies (TWAS), TWAS fine-mapping^51^, eQTL fine-mapping, estimation of *trans*-heritability, estimation of environmental effects, identifying disease-critical gene sets, tissues, and gene-cell-type pairs^52,53,54^, estimating SNP-disease effects mediated by changes in gene expression^55^, and partitioning gene expression heritability by cell types or functional categories using stratified LD score regression^56^ – and naturally, application of these methods to scRNA-seq data will only become more frequent. Therefore, inaccuracies in *cis*-heritability estimation resulting from pseudobulking can substantially distort the interpretations of biological mechanisms and disease pathology. However, no strategies currently exist to estimate the *cis*-heritability of gene expression using native scRNA-seq data.

Here, we propose a generalized Poisson mixed effects model called scGeneHE (single cell Gene expression Heritability Estimation) that leverages both inter-individual (relatedness among people) and intra-individual (relatedness among cells of the same person) variation in scRNA-seq expression measurements to improve the accuracy of gene expression *cis*-heritability estimation and error rate of eGene detection. scGeneHE addresses the high sparsity rate and discrete read counts native to scRNA-seq data by employing a Poisson distribution to model gene expression across cells and donors. Our framework shares the same Poisson mixed effect model of the recent single cell eQTL mapping method SAIGE-QTL^49^, which overcomes the inherent computational inefficiency of modeling thousands of cells and their variance-covariance matrices for mixed-effects models by expressly avoiding matrix inversions thanks to the use of preconditioned conjugate gradients. Notably, we estimated that employing the widely-used GCTA heritability estimation framework (with an additional customized random effect for intra-individual variation) would be intractable when applied to millions of cellular measurements across tens of thousands of genes, in addition to being naturally underpowered due to its assumption of normally distributed count data. scGeneHE decomposes the variance of gene expression measurements into genetic (inter-individual) and shared cellular environment (intra-individual) variances. The latter is especially critical, as the shared environment of cells originating from the same donor is the leading cause of overestimation and miscalibration in pseudobulk heritability analysis. In simulations, scGeneHE has a controlled false positive rate at or below 5% and achieves less biased estimates than the pseudobulk approach, at the expense of lower power at smaller *cis*-heritabilities, but comparable power at higher *cis*-heritabilities. We applied scGeneHE to data from the OneK1K cohort, the largest available population-scale scRNA-seq dataset with paired genotyping data^27^. Our results unveil cell-type-specificity of regulatory mechanisms for hundreds of genes related to immune-mediated diseases. Overall, scGeneHE demonstrates its efficiency and robustness in identifying causal genetic regulatory components in both simulated and real data analysis.

## Results

### Overview of Method

scGeneHE estimates the gene expression heritability explained by the *cis*-genetic component of gene expression within a cell type using the resolution of single-cell gene expression profiles. To address the sparsity and repeated measurements in scRNA-seq data, scGeneHE employs a Poisson linear mixed model for heritability estimation. The model directly captures gene expression counts through a log-linear transformation, associating them to fixed effect covariates and random effects while ensuring positive parameter values, e.g., variance explained by shared genetics and variance explained by shared environment. Specifically, the model integrates cell-level covariates, such as gene expression principal components (PCs), mitochondrial content, and total normalized unique molecular identifier (UMI) counts, along with individual-level covariates, including genotype principal components, demographic factors such as age and sex, and batch effects. These covariates are treated as fixed effects, while the model accounts for random effects that capture genetic relatedness among individuals and cell-cell relatedness within individuals (**Figure 1C**). These random effects are assumed to follow multivariate normal distributions, with covariance structures informed by genetic and cell-cell relationships (**Methods**). Since Poisson distributions are defined without (environmental) noise, we use the cell-cell relatedness random effect term to capture all sources of non-genetic variation caused by repeated measurements per individual. scGeneHE employs the individual-level genetic relationship matrix (GRM) using *cis*-SNPs (within +/-500 kilobases of the gene) and expands it to the level of single cells via row-wise duplication. For simplicity, we construct the cell-cell relatedness matrix (CRM) by assuming that cells of a particular cell type within each person are perfectly related to each other (covariance of 1) due to sharing the same environment and are not related at all to cells from different individuals (covariance of 0), beyond sharing the same *cis*-genetic component of gene expression, e.g., heritability.

scGeneHE simultaneously estimates variances of the genetic component of gene expression and the non-genetic cellular relatedness using a penalized quasi-likelihood as previously employed in single-cell eQTL analysis^49^. Heritability (*h*^*2*^_*ge*_) is calculated as the proportion of variance attributable to genetic factors, constrained to the range [0,1] to be comparable to Gaussian heritability values^57^. Parameters are optimized using the average information restricted maximum likelihood, iteratively refining estimates of fixed effects, random effects, and variance components, as similarly employed in GCTA^3^. To determine statistical significance, we use a non-parametric bootstrap approach to estimate the standard error of heritability, followed by a one-sided t-test to evaluate the likelihood of obtaining the same estimate if the null hypothesis (*h*^*2*^_*ge*_ = 0) was true. Genes with significantly non-zero 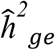 are deemed *cis*-heritable, also referred to as eGenes below.

### Simulations comparing cis-heritability estimation with pseudobulk counts to scGeneHE

We conducted extensive simulations to evaluate the performance of scGeneHE under various genetic architectures using real genotypes. Focusing on a 1 megabase (Mb) region of chromosome 1, we simulated gene expression for European individuals from the 1000 Genomes Project^58^. For each individual, single-cell gene expression was simulated by combining a shared genetic component, determined by predefined *cis*-heritability (*h*^*2*^_*ge*_), with random environmental noise (**Methods**). Causal *cis*-eQTL effect sizes were sampled from normal distributions with mean 0 and variance equal to the per-SNP *cis*-heritability, e.g., total *h*^*2*^_*ge*_ divided by the number of *cis*-eQTLs per gene (default is 5); non-causal SNPs were assigned effect sizes of zero. To mimic technical dropout, sparsity was introduced by setting a percentage of cell-level gene expression profiles to zero^59^. Null analyses were performed by permuting donor-cell assignments, thereby disrupting genotype-expression associations. Pseudobulk gene expression was simulated by averaging or summing cell-level data within individuals, where both approaches generated the same *cis*-heritability estimates; therefore, we only consider results from summing cell-level read counts below (**Supplementary Figure 1**). Our simulation model incorporates the top six genotyping PCs and randomly simulated mitochondrial gene percentages as covariates. Heritability estimation for *cis*-heritable and null genes was performed using scGeneHE with bootstrap resampling to compute standard errors. Significance was assessed with a one-sided t-test using a nominal significance threshold of p < 0.05; power and type I error rates were calculated for both single-cell and pseudobulk analyses. For pseudobulk data, heritability and corresponding standard errors were estimated using GCTA^3^. We systematically varied simulation parameters to examine their effects on *cis*-heritability estimates, including changes in true *cis*-heritability, donor sample size, number of cells per individual, number of causal eQTLs, and sparsity rates. These simulations provide a comprehensive framework for inferring how scGeneHE might perform in realistic scenarios. Please see **Methods** for a more detailed overview of our simulation framework.

We first evaluated the bias of scGeneHE estimates of heritability on both causal (genes with a non-zero value of *cis*-heritability) and null (not *cis*-heritable) genes. scGeneHE generates unbiased estimates of gene expression heritability across a range of true heritability levels (1%, 5%, 10%, 25%, and 50%) for causal genes, where the true heritability is captured within the 95% confidence interval of estimates across independent simulations (**Figure 2A, Supplementary Table 1**). In comparison, the widely used approach of applying GCTA to pseudobulk gene expression profiles exhibits a systematic upward bias at larger values of true *cis*-heritability and great uncertainty at smaller values of true *cis*-heritability. For null genes, scGeneHE consistently reports near-zero heritability with slightly smaller confidence intervals on average compared to the pseudobulk approach, although the latter does not estimate significantly non-zero estimates of *cis*-heritability on average. We note that while scGeneHE can only estimate non-negative parameter values, the distribution of null gene estimated *cis*-heritabilities is not significantly greater than zero at any value of true *cis*-heritability. Overall, by effectively modeling cell-level variation and accounting for the sparsity and heterogeneity inherent in scRNA-seq data, scGeneHE reduces bias on causal estimates of *cis*-heritability relative to the pseudobulk approach.

**Figure 2.**
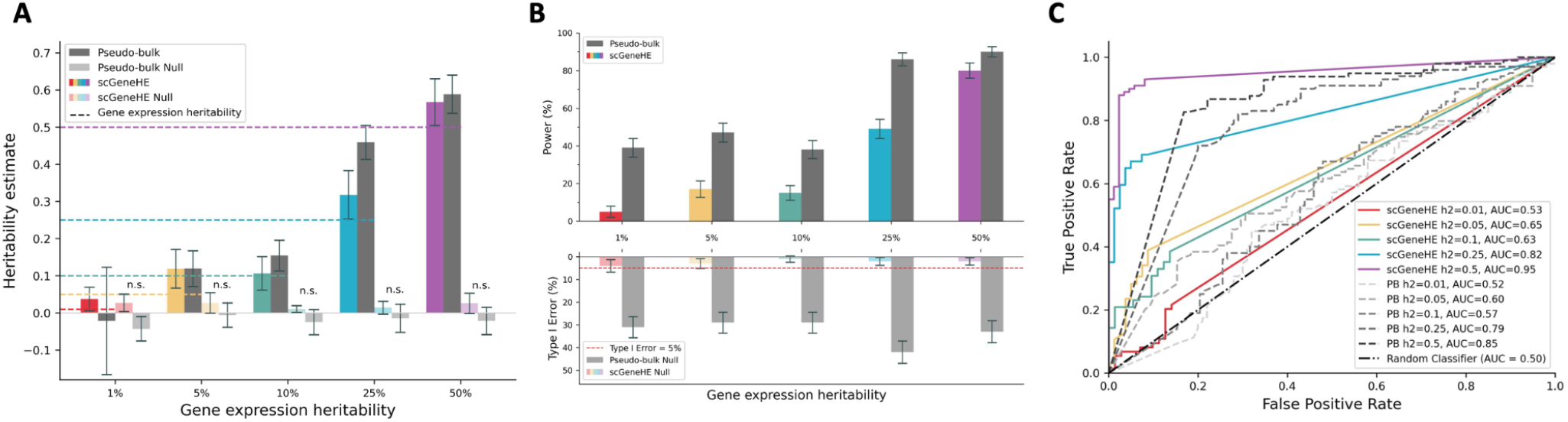
Simulations reveal improved calibration of scGeneHE compared to pseudobulk analysis of scRNA-seq data. (**a**) Estimation of causal gene expression heritability 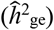 by scGeneHE (color) relative to pseudobulk analysis (gray). Null calibration is assessed via a permutation analysis in which gene expression values across cells and donors no longer correspond to genotypes: scGeneHE null calibration indicated by lightly colored bars, pseudobulk null calibration indicated by light gray bars. Colorful dotted lines indicate true value of gene expression heritability 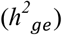 indicated on the x-axis. Bar height represents the average estimate of gene expression heritability 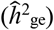 across 100 independently simulated genes. Black line segments represent the 95% confidence intervals, using the standard deviation across independent simulations. (**b**) Power and type I error rate of scGeneHE and pseudobulk analysis. Bar height represents the average proportion of 100 independently simulated genes for which scGeneHE successfully estimated a non-zero value for 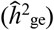 in causal simulations (light blue) or null simulations resulting from permutation analysis. Power and type I error is similarly depicted for pseudobulk analysis with light gray (causal) and dark gray (null) bars. Black line segments represent the 95% confidence intervals, using the standard deviation across independent simulations. (**c**) Area under the receive operator characteristic curve (AUROC) where the true positive rate (or sensitivity) is on the y-axis and the false positive rate (or 1 - specificity) is shown on the x-axis. 1000 uniformly spaced p-values were used to generate this plot. For all panels, 100 donors were modeled, true gene expression heritability was varied between 1% and 50%, each gene had 5 causal eQTLs, and for each donor, 50 cells were simulated where a sparsity rate of 80% is applied. Numerical results are reported in **Supplementary Tables 1-3**.

We next evaluated the power and type I error of scGeneHE on both causal and null genes, respectively (**Figure 2B, Supplementary Table 2**). While previous studies have employed the pseudobulk approach to identify *cis*-heritable genes^45,47^, it is not surprising that this approach has relatively high power. The power of scGeneHE is modest in comparison, although this effect is less prominent at larger heritabilities. However, the discrepancy in power is moot because the pseudobulk approach is susceptible to large type I error rates (~30% or greater across different true *cis*-heritabilities), while scGeneHE consistently maintains a type I error rate less than 5%. Therefore, in real data analysis, one may expect to have little issue detecting a truly *cis*-heritable gene; however, distinguishing such a gene from false positives will be very challenging by using pseudobulk approach. We then utilized ROC (receiver operator characteristic) curves to simultaneously compare the tradeoffs between power and type I error between the two approaches (**Figure 2C, Supplementary Table 3**); for example, compare the purple line with the dashed black line. scGeneHE consistently achieved higher AUROC (area under the ROC) values across all tested values of *cis*-heritability, with the greatest improvements at larger *cis*-heritabilities: when *h*^*2*^_*ge*_ was set to 0.5, scGeneHE obtained an AUROC of 95% while pseudobulk obtained 85%. Overall, our analysis reveals the shortcomings of *cis*-heritability estimation using the pseudobulk approach, originating from its treatment of highly correlated gene expression profiles as independent measurements, resulting in artificially boosted estimates of *h*^*2*^_*ge*_ and artificially smaller standard errors on the estimate. To further investigate the latter statement, we found that the distribution of bootstrap standard errors produced by scGeneHE is centered as expected around the empirical standard deviation of estimates of *h*^*2*^_*ge*_ across independent simulations (**Supplementary Figure 2**). However, this is not the case for the pseudobulk approach, in which the distribution of standard errors produced by GCTA is smaller than one would expect given the empirical standard deviation of estimates of *h*^*2*^_*ge*_ across independent simulations. Overall, by modeling dependencies among gene expression profiles from each cell, the scGeneHE framework presents a well-calibrated and modestly powerful method to estimate the *cis*-heritability of gene expression from single cell RNA-sequencing data.

## The effect of variable shared cellular environment on cis-heritability estimation

As noted above, the generalized mixed model of scGeneHE makes a simplifying assumption regarding the perfectly shared environmental component of cells of a given cell type that come from the same individual. As a result, scGeneHE employs an identity matrix as the correlation matrix of cell-cell relatedness. We note that this is not designed to capture the substructure of cell types shared across individuals, but rather to capture substructure simply within an individual, which may be unique to that individual. We next sought to assess how this simplifying assumption would affect *cis*-heritability estimation when more complex cell-cell relatedness is present in the data. We thus create a framework to simulate various environmental cellular heterogeneity (**Figure 3A**). When simulating gene expression, the *cis*-genetic component is identically shared among cells within an individual. In this section, we introduce an index of cellular heterogeneity *K* when generating environmental noise. All cells within an individual are divided into *K* groups where each group of cells share the same environmental noise, but the environmental noise of cells from different groups are independently sampled from the same normal distribution with variance controlled by the desired true *cis*-heritability. A smaller value of *K* represents greater similarity in cellular environment, analogous to a fine-grained clustering of cells with similar functionality and origin. In reality, we anticipate that the lowest values of *K* are not realistic, as even cells from the same functional cell type will have different environmental influences due to differences in their spatial position, cellular neighbors, and resulting cell-cell signaling. A larger value of *K* represents greater heterogeneity in the cellular environment; in the most extreme case, when *K* equals the number of cells, this represents each cell serving a distinct biological function or coming from a distinct origin, such as may be the case if no cell type clustering was performed on raw scRNA-seq data. Since we later apply scGeneHE to real gene expression profiles of cell types that have been identified from unsupervised clustering algorithms and similarity metrics, intermediate or small values of *K* are expected to be most relevant to assessing model performance.

**Figure 3.**
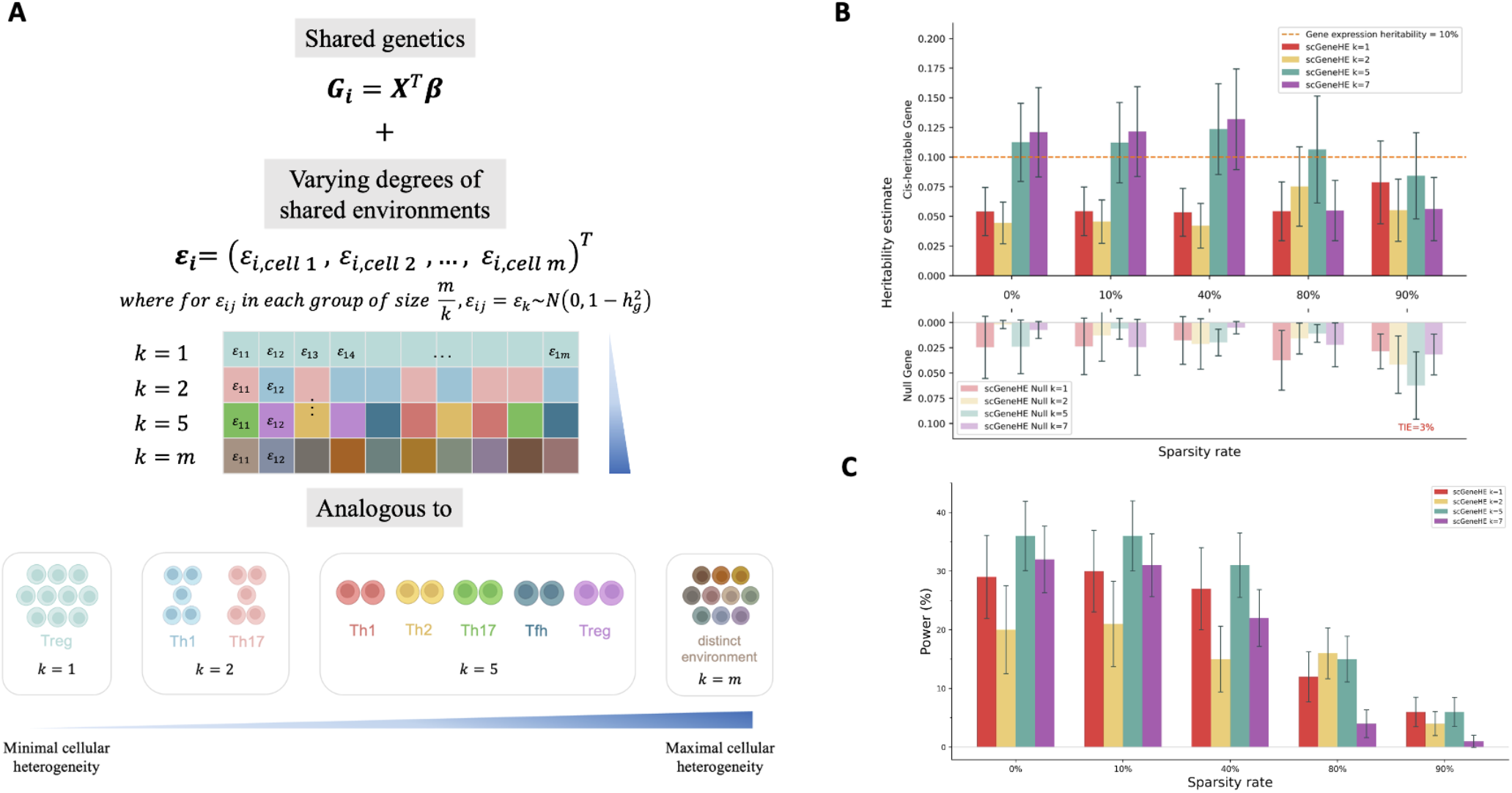
Modeling heterogeneity in cellular environments. (**a**) Schematic depicting the choice of different values of K (number of unique cellular environments from which cells of a given cell type are derived). K = 1 indicates that all cells come from an identical cellular environment and thus their gene expression levels in a donor are identical (before sparsity is applied). K = m indicates that each cell comes from a unique cellular environment, where the variance of this heterogeneity is equal to 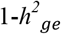. (**b, top**) scGeneHE *cis*-heritability estimates for different values of K across various levels of sparsity. (**b, bottom**) Inverted bar graph depicting the *cis*-heritability estimates for genes with no true genetic component, derived from permutation analysis. The largest bias is observed for K = 5 and 90% sparsity; therefore, we annotate the type I error rate to demonstrate well-maintained calibration of scGeneHE. (**c**) scGeneHE power to detect eGenes across different values of K and sparsity. In all panels, black line segments represent the 95% confidence interval calculated using the standard deviation across 100 independent simulations. Numerical results are reported in **Supplementary Tables 4-5**.

Next, we evaluated the bias of scGeneHE estimates across varying levels of cellular sparsity (0%, 10%, 40%, 80%, and 90%) while varying cellular heterogeneity (*K* =1, 2, 5, 7) for both causal and null genes. We note that in previous analyses, we set *K* = 5 (**Figure 2**). For the *cis*-heritable genes, scGeneHE generates results approaching the true *cis*-heritability without overestimation (**Figure 3B, Supplementary Table 4**). While scGeneHE underestimates the true *cis*-heritability by about half when cellular heterogeneity is low (*K* < 3), this trend is expected because the inter-individual genetic variation is confounded by (e.g., is highly correlated with) the intra-individual (cellular) variation across gene expression profiles. However, we continue to note that such low values of *K* are not physiologically likely. As a result, scGeneHE mistakenly attributes about half of the *cis*-heritability to shared cellular environment; this trend also occurs even for larger values of *K* at greater percentages of sparsity for similar reasons. For null genes, scGeneHE shows slightly inflated estimates of heritability for larger levels of sparsity but effectively controls the type I error below the nominal threshold of 5% even for the setting resulting in the largest upward biased estimate of heritability (*K* = 5, sparsity = 90%, type I error = 3%). This highlights scGeneHE’s ability to differentiate *cis*-heritable from null genes, even under conditions of high sparsity and/or environmental heterogeneity.

We also tested the performance of scGeneHE in secondary simulations by separately varying the number of donors, the number of causal *cis*-eQTLs, and the number of cells per donor. First, we found that the scenario with largest donor sample size (*n* = 500) and moderately heterogeneous cellular environments result in inflated *cis*-heritability estimates, however the type I error is still well-controlled (**Supplementary Figure 3**). This is not the case for pseudobulk analysis where we observe both overestimation and exceedingly large type I error. Second, we varied the number of causal *cis*-eQTLs and found that when there are fewer causal eQTLs and also a moderately heterogeneous cellular environment, scGeneHE overestimates heritability – but again, the type I error is relatively well-controlled (**Supplementary Figure 4**). This is again not the case for the pseudobulk approach which experiences similar trends of overestimation but is coupled with exceedingly high type I error. Lastly, we found that varying the number of cells per donor minimally affects the performance of scGeneHE (**Supplementary Figure 5**).

Lastly, we considered the power to detect *cis*-heritable genes across different sparsity and environmental heterogeneity levels (**Figure 3C, Supplementary Table 5**). We observed that performing null analyses at high sparsity is not completely effective, as the multitude of zeros creates its own structure in the data causing the false perception of a shared genetic component. Across 100 independent simulations, we observe that power decreases consistently with increasing sparsity. On the other hand, the value of *K* does not result in a consistent effect on eGene detection power, except for both large *K* and large sparsity (80-90%). Overall, even with the simplifying assumptions we make while modeling cell-cell relatedness, scGeneHE effectively handles varying levels of cellular sparsity and heterogeneity, avoiding overestimations of *cis*-heritability while maintaining robust type I error control for null genes. We conducted similar analyses for the pseudobulk approach, which show greater overestimates of *cis*-heritability for larger values of *K* and at most sparsity rates (**Supplementary Figure 6**) and high power to detect true eGenes (**Supplementary Figure 7**), which remains misleading given its inflated false positive rates in our primary simulations. For null genes, again consistent with our primary simulations, the pseudobulk approach produces *cis*-heritability estimates mostly centered around zero (**Supplementary Figure 6**); although, we note that small standard errors in this regime still result in large false positive rates on eGene detection. Overall, we find that our simplifying assumption of using a block diagonal for cell-cell relatedness is sufficient to properly estimate *cis*-heritability of gene expression, even in the presence of cell type substructure.

## Identifying cell-type specific gene regulation critical to immune-mediated disease

We applied scGeneHE to the OneK1K RNA-sequencing cohort of 969 individuals and 822,552 cells across 11 immune cell types^27,60^. In order to increase the computational feasibility of our analysis and not compromise statistical power, we selected 11 cell types for which there were between 10 and 90 cells per donor; this excluded 13 cell types with insufficient cell counts and 3 cell types with exceedingly large numbers of cells per donor: natural killer cells, CD8+ T effector memory, and CD4+ T central memory cells. Given the breadth of immune cell types and finely resolute subpopulations of T cell types in our dataset, we first identify 1,211 genes that have been mapped to previously identified genome-wide significant loci for rheumatoid arthritis (RA) and other immune-mediated diseases and traits (**Supplementary Table 6**)^61^. We additionally required that these genes be expressed in >10% of cells (to ensure estimation convergence) from at least 2 analyzed cell types, to ensure that we would have sufficient expression data to estimate *cis*-heritability with scGeneHE, resulting in a total of 166 genes. We set out to compare scGeneHE *cis*-heritability estimates with pseudobulk and bulk approaches. To this end, we generated pseudobulk expression profiles by summing the normalized TPM (transcripts per million) read counts across cells belonging to the same individual followed by log-scaling (**Methods**) and consulted our previously estimated bulk-level *cis*-heritabilities for genes that are expressed in GTEx whole blood tissue^18,53^. In both our scGeneHE and pseudobulk heritability analyses, we included the following covariates: sex, age, pool for batch effect, mitochondrial gene percentage, log-transformed UMI count per cell for single-cell analysis or summed linear-scale UMI count for pseudobulk analysis, 6 genotype PCs, and 2 cell-type-specific gene expression PCs, as previously employed^27^. For bulk analysis, we included five genotyping PCs, gene expression PEER (probabilistic estimation of expression residuals) factors from GTEx, and sequencing platform, as previously described^53^.

We then compared the detection of eGenes and the value of *cis*-heritability estimates for all pairs of genes and cell types across various resolutions of gene expression profiles: single-cell, pseudobulk, and bulk. We performed multiple hypothesis correction across the number of cell types tested per gene to properly control the false positive rate, under the assumption that each gene has a true *cis*-genetic component in at least one cell type. Of the 166 genes tested, we identified 25 genes that have a significantly non-zero *cis*-heritability estimate according to scGeneHE and 88 genes that have a significantly non-zero *cis*-heritability estimate according to pseudobulk analysis. Given the inflated type I error and overestimation of *cis*-heritability by the pseudobulk approach in our simulations, we anticipated that pseudobulk analysis would detect more eGenes than scGeneHE; indeed we observed these 63 genes for which scGeneHE could not recapitulate the detection of a *cis*-genetic component indicated by the pseudobulk approach in any cell type. The 25 genes that were detected as an eGene in at least one cell type by scGeneHE are displayed in **Figure 4** (**Supplementary Table 7**).

**Figure 4.**
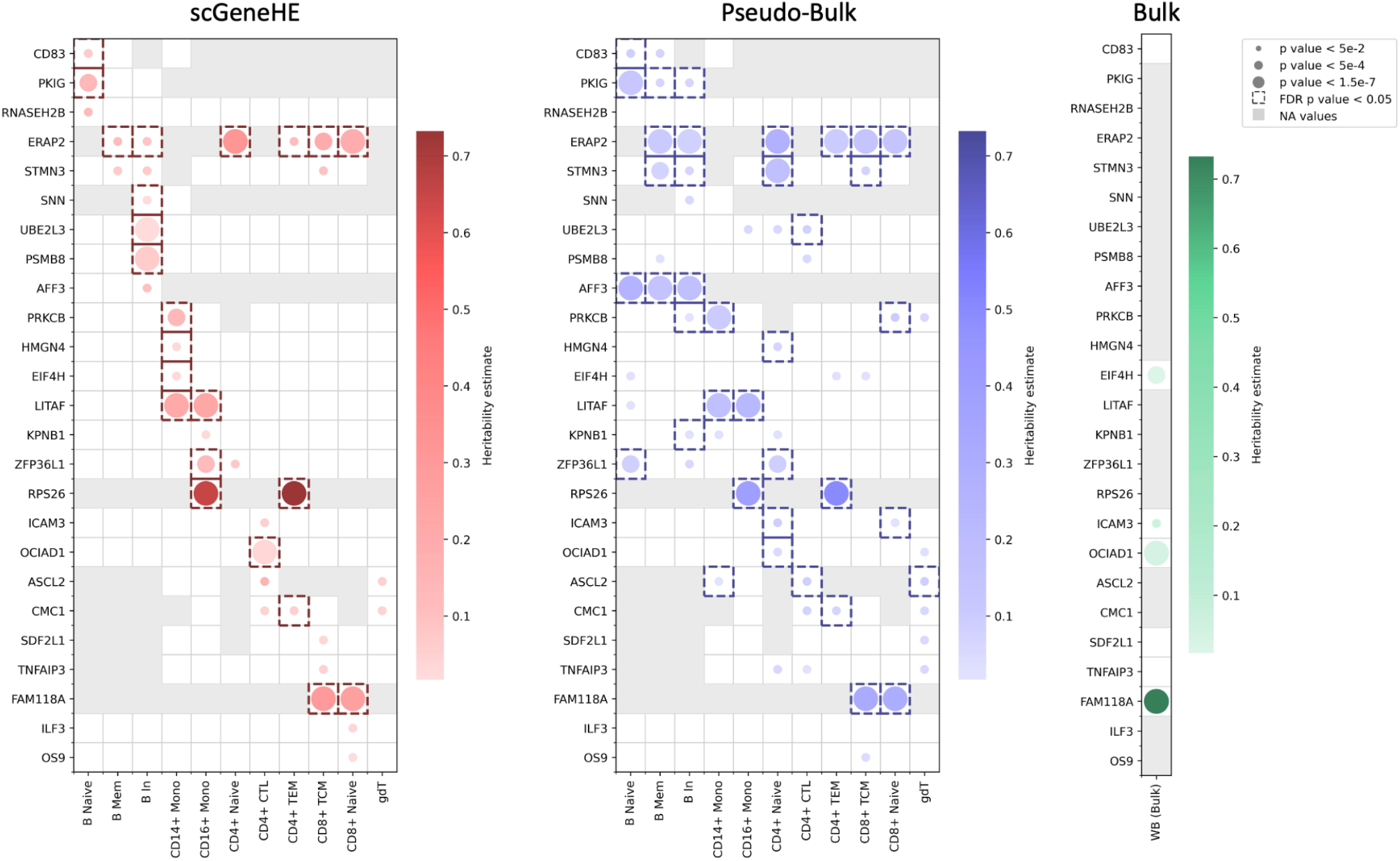
*Cis*-heritability estimates in OneK1K single cell-RNA sequencing data. (**a**) *Cis*-heritability estimates resulting from applying scGeneHE to single cell gene expression profiles. (**b**) *Cis*-heritability estimates resulting from applying GCTA to pseudobulk gene expression values. (**c**) *Cis*-heritability estimates from bulk whole blood tissue in GTEx using GCTA. For all panels, gray boxes indicate that the gene was too lowly expressed in a given cell type (< 10% of cells) (or in bulk analysis, the gene was not present) and thus we did not attempt to estimate *cis*-heritability. The color of dots refers to the magnitude of heritability estimated, the size of dots is inversely proportional to the nominal p-value of the heritability estimate, and dashed boxes indicate that heritability was significantly non-zero at 5% FDR (false discovery rate) after correcting for the various cell types tested for that gene. B mem, B memory cells; B In, B intermediate cells; Mono, monocytes; CTL, cytotoxic T lymphocytes; TEM, T effector memory cells; TCM, T central memory cells; gdT, gamma-delta T cells; WB, whole blood. Numerical results are reported in **Supplementary Table 7**.

Of these 25 and 88 genes, 7 and 13 genes, respectively, had a significantly non-zero *cis*-heritability estimate in whole blood bulk tissue analysis. In general, the GTEx bulk heritability estimates from whole blood poorly capture the cell-type-specificity of gene regulation in OneK1K cell types and will be relatively underpowered to detect eGenes, especially when these eGenes are specifically regulated in cell types that make up only a small proportion of whole blood. Amongst genes that are detected as *cis*-heritable in GTEx whole blood tissue, scGeneHE can help pinpoint the specific cell type(s) harboring the *cis*-genetic component of gene expression, such as identifying that *ICAM3* is specifically genetically regulated in CD4+ cytotoxic lymphocytes (CTLs). Indeed, *ICAM3* is critical in initiating the immune response of T lymphocytes by regulating lymphocyte morphology and the ability to bind to antigen-presenting cells^62,63^. Based on this previous work, results from both scGeneHE and the pseudobulk approach (implicating CD4+ and CD8+ naive T cells) seem biologically plausible. Because the bulk analysis is able to detect this eGene, its *cis*-eQTL effects may be very large in CD4+ CTLs or this gene may have a *cis*-genetic component that is shared with other constituent cell types of whole blood that are not analyzed here.

Next, we delve deeper into some notable differences in results between the scGeneHE and pseudobulk approaches. First, despite the increased power (albeit large false positive rate) of the pseudobulk approach observed in simulations, there are a few genes (*RNASEH2B* and *ILF3*) for which scGeneHE detected a significant *cis*-heritable component of expression, which the pseudobulk approach could not. These findings are broadly supported by previous studies. For instance, *RNASEH2B* has been implicated in immune-related processes and systemic autoimmunity, including a role in the pathogenesis of the auto-inflammatory Aicardi-Goutieres syndrome via dysregulated nucleic acid metabolism^64^. Moreover, its cell-type-specificity in B naive cells seems biologically plausible due to increased R loop activity in resting B cells, which could lead to cellular stress and enhanced autoimmune response^65^. scGeneHE also identified that *ILF3* is genetically regulated specifically in CD8+ naive T cells, which has been shown to be a possible target for rheumatoid arthritis treatment by regulating IL-2 expression in T cells and thus promoting synoviolin expression in synovial cells^66^. Separately, cancer immunotherapy studies have shown that disruption of the ILF3 protein complex results in reduced CD8+ T cell infiltration^67^.

Secondly, we hypothesized that scGeneHE would refine the generic, cell-type-nonspecific *cis*-heritability estimates made by the pseudobulk approach to specific cell types. For 72% of genes, pseudobulk analysis estimated non-zero *cis*-heritability in > 1 cell type and 67% of the time, scGeneHE refined these multiple signals to a single cell type. For instance, scGeneHE detects B cell specific genetic regulation of *CD83, PKIG*, and *AFF3*, and CD14+ monocyte specific regulation of *PRKCB*, an exemplar gene whose *cis*-heritability is discussed in the next subsection. This is consistent with the established roles of these genes in immune signaling and lymphocyte activation: *CD83* plays a critical role in naive B cells by initiating the dynamic progression of B cell activation in rheumatoid synovial ectopic lymphoid structures^68^. Moreover, we note that *CD83* was not detected as *cis*-heritable in bulk whole blood tissue, supporting the likelihood of its cell-type-specificity; B cells represent only a trace fraction of whole blood tissue (only 1% of whole blood tissue are lymphocytes, of which B cells represent a small percentage). For *PKIG*, scGeneHE refined the generic B cell signature from pseudobulk analysis to specifically naive B cells, supported by a recent study showing that *PKIG* expression levels affect acute myocardial infarction severity dependent on the relative abundance of naive B cells^69^. For *AFF3*, one of the original genes implicated by early RA GWAS^70^, scGeneHE estimated that *cis*-genetic regulation is localized to intermediate B cells, which is consistent with *AFF3*’s primary role in B cell development and immunoglobulin class switch recombination mechanisms studied with murine models of non-naive B cells^71^. Lastly, we also observed while the pseudobulk *cis*-heritability of *ERAP2* was found to be consistent large and highly significant across many cell types, results from scGeneHE were much more variable, with CD4+ naive T cells accounting for the strongest significance and highest *cis*-heritability. Consistent with our estimation, *ERAP2* has been shown to have a dual role in autoimmune diseases by influencing antigen presentation to T cells while also providing protective effects against severe infections^72^.

Lastly, we focus on examples where scGeneHE and the pseudobulk approach identify entirely different cell types harboring *cis*-genetic regulatory mechanisms. First, for example, scGeneHE detects cell-type-specific CD16+ monocyte heritability for *ZFP36L1*, while the pseudobulk approach claims specificity in naive B cells. However, *ZFP36L1* has an established role in positively regulating monocyte and macrophage differentiation while also contributing to anti-inflammatory processes in autoimmune diseases such as rheumatoid arthritis, whereas B cells are more associated with pro-inflammatory processes^73^. We discuss the plausibility of this gene-cell type pair in more depth in the next subsection. Second, for *HMGN4*, scGeneHE implicates CD14+ monocytes while pseudobulk implicates naive CD4+ T cells. *HMGN4* is known to play a key role in oncogenesis mediated by STAT3, which has been implicated in the transition of cell fate between CD14+ and CD16+ monocytes^74,75^. Third, scGeneHE identifies biologically plausible cell-type-specific regulation of *UBE2L3* in intermediate B cells, whereas pseudobulk detects *UBE2L3* as an eGene for CD4+ cytotoxic lymphocytes. Prior work has established that *UBE2L3* is associated with increased susceptibility to systemic lupus erythematosus, where *UBE2L3* promotes higher autoimmune activity and elicits responses from abnormal CD19+ B cells, a type of intermediate B cell^76^. Fourth, scGeneHE implicates CD8+ T central memory (TCM) cells in the *cis*-regulation of *TNFAIP3*. Although *TNFAIP3* contains a fine-mapped risk variant for RA and type 1 diabetes, this does not necessarily imply that the risk variant is an eQTL of *TNFAIP3*^*77*^. Interestingly, previous work supports that *TNFAIP3* inhibits CD8+ T cells from performing their natural antitumor activity and downregulation of *TNFAIP3* may be helpful in promoting tumor rejection by the immune system^78^. We also note that the *cis*-genetic component of *TNFAIP3* is likely highly specific to lymphocytes, as bulk analysis did not detect this gene as an eGene.

Overall, scGeneHE helps identify the *cis*-genetic component of gene expression in scenarios where the pseudobulk approach was underpowered or was cell-type-nonspecific. We also use scGeneHE to contest the cell-type-specific *cis*-heritability estimates of the pseudobulk approach. As a result, our method can help clarify the cell-type-specific localization of eQTL and disease-critical variant effects.

### Traversing cis-heritability of gene expression through varying granularity of immune cell subtypes

In scRNA-seq data analysis, a mix of unsupervised and supervised clustering is often performed to group cells by similar gene expression profiles and thus cellular functionality. The *cis*-genetic component of gene expression is likely to be most consistent and most powerfully detected in a fine-grained cellular population, as there is minimal variation in gene expression profiles and possibly true eQTL effects. However, additionally estimating *cis*-heritability in less fine-grained cellular populations, e.g., with more functional heterogeneity, could reveal shared gene regulatory effects in constituent cell types. Thus, we set out to investigate which genes have patterns of *cis*-genetic regulation that are more broadly detected in less differentiated cell types in a cell lineage versus those that are only specific to highly differentiated cell types. To this end, we focus on the myeloid lineage within our scRNA-seq dataset. **Figure 5A** demonstrates our progressive clustering of immune cells, starting with peripheral blood mononuclear cells (PBMCs) as the most inclusive, heterogeneous population, followed by the increasingly more fine-grained cellular populations: myeloid cells, monocytes, and the two monocyte subsets: CD14+ and CD16+ monocytes. To ensure consistent power to identify *cis*-heritable genes, we downsampled all cell type clusters to the same donor size (**Methods**). We relate this analysis back to **Figure 3A**, where our analysis of PBMCs may be better anticipated by simulations with a larger value of *K*, while our analysis of CD14+ and CD16+ monocytes may be better anticipated by simulations with a smaller value of *K*. This framework allows us to assess trajectories of *cis*-heritability across cellular populations with varying similarity in expression profiles.

To this end, we applied scGeneHE to 200 randomly selected genes that were expressed in at least 10% of cells and estimated *cis*-heritability across PBMCs, myeloid cells, monocytes, CD14+ monocytes, and CD16+ monocytes (**Figure 5B, Supplementary Table 8**). We first note that several genes were detected as *cis*-heritable in PBMCs, but were not found to be *cis*-heritable in any myeloid lineage cell types and thus could be involved in the other immune lineages present in our dataset. We hypothesize that such genes will often be involved in more generic and fundamental cellular processes; for instance, the genes *MZT2B* and *RHOC* are involved in cytoskeletal organization and may be less likely to exhibit high *cis*-heritability in more fine-grained cell types^79,80^. For genes that were found to be nominally *cis*-heritable in PBMCs and myeloid cells, a similar explanation may be given – either the genetic regulation of the gene is to control a fundamental cellular process or to control a dendritic cell-specific process, since we did not evaluate this specific lineage. In the case of the ubiquitously expressed *EIF4B* gene, which initiates protein translation according to the HUGO gene nomenclature committee, the former hypothesis regarding fundamental functionality is more likely^81^. For genes with nominal non-zero *cis*-heritability in the most heterogeneous cell types but zero *cis*-heritability in the most homogeneous cell types (such as RPA3), we hypothesize that there may be mutually exclusive true eQTL effects in CD14+ and CD16+ monocytes that are individually small. Therefore, it is difficult for scGeneHE to classify these genes as eGenes; however, in the more heterogeneous cell types, scGeneHE may be able to aggregate these mutually exclusive effects to increase power to detect the gene as an eGene.

**Figure 5.**
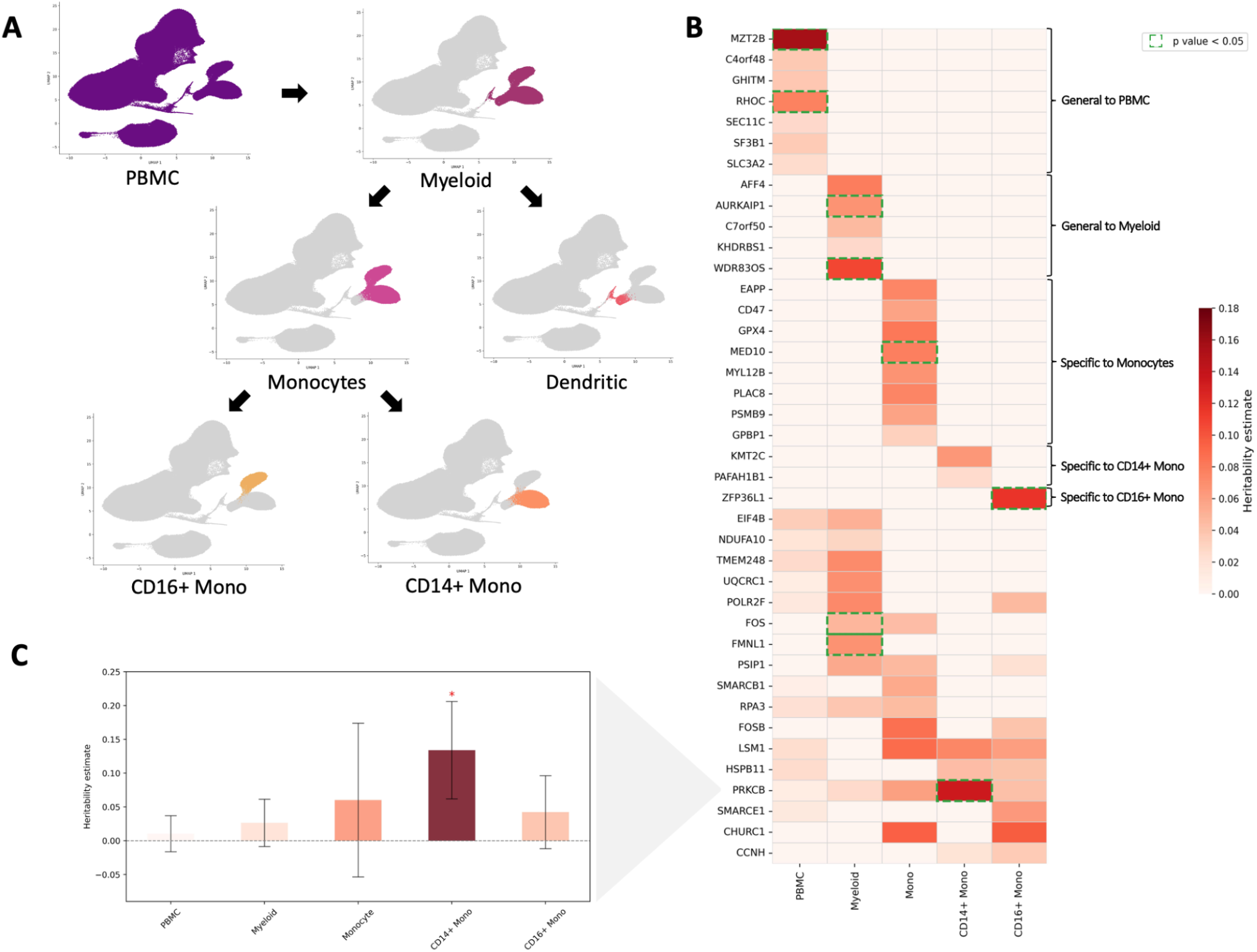
*Cis*-heritability estimates through varying granularities of immune cell subtypes. (**a**) Relation between myeloid lineage cell types: PBMCs are the most heterogeneous cell type, of which a fraction are myeloid cells, myeloid cells can be partitioned into monocytes and dendritic cells; monocytes can be further partitioned into CD16+ and CD14+ monocytes. (**b**) *Cis*-heritability estimates resulting from applying scGeneHE to single cell gene expression profiles. Genes that are significant in specific cell types after multiple hypothesis correction are outlined by green dashed lines. (**c**) Example gene *PRKCB* exhibits increasing 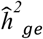 as immune cell types are more resolutely defined, reaching statistical significance in CD14+ monocytes (red asterisk). PBMC, peripheral blood mononuclear cells; Mono, monocytes. Numerical results are reported in **Supplementary Table 8**.

We observed another pattern of *cis*-heritability in which genes were detected as *cis*-heritable in both monocytes and one specific monocyte subtype, such as *CHURC1*. We hypothesize that in this case, the true effect sizes of the eQTLs in CD16+ monocytes were large enough to be detected in monocytes, despite this cell type being a mixture of CD14+ and CD16+ monocytes. *CHURC1* is a gene involved in the fibroblast growth factor receptor (FGFR) signaling pathway according to NCBI and the FGFR pathway has recently been shown to play a role in the specified differentiation of monocytes and macrophages (also referred to as tissue-resident monocytes)^82,83,84^.

On the other extreme, we explore some exemplar genes where the signal to noise ratio for estimating heritability was overcome only by considering the most specific and functionally similar monocyte subtypes. In these scenarios, we hypothesize that eQTL effects are relatively small and specific to a subset of monocytes, and thus not able to be detected when that subset is just a fraction of a larger cellular population such as monocytes or myeloid cells. *ZFP36L1* was detected as *cis*-heritable only in the most specific subtype of myeloid cells – CD16+ monocytes. Notably, for this gene, all other less differentiated cell types produced a *cis*-heritability estimate of zero. Previous work has demonstrated that this gene interacts with *CDK6*, which promotes monocyte differentiation and maturation into specific subtypes, such as CD16+ monocytes^73,85^. The *HSPB11* gene exhibits similar specificity, although not maintaining significance after correction for the number of cell types tested, but has not previously been implicated in monocyte regulation. Regarding specificity for the other monocyte subset, *PRKCB* was detected as significantly *cis*-heritable only in CD14+ monocytes. This result is consistent with a recent study that mapped two genome-wide significant variants associated with nontuberculous mycobacterial infection to *PRKCB*, for which scQTLbase reports eQTL associations with CD14+ monocytes rather than other monocyte populations^86,87^. Interestingly, tracing the *cis*-heritability estimates for *PRKCB* from less to more differentiated populations within the myeloid lineage shows increasing estimates that require the specificity of the CD14+ subpopulation to reach statistical significance **(Figure 5C)**. We note that this result is robust to downsampling cells even to 20% of the original amount **(Supplementary Figure 8)**. Although not maintaining significance after multiple hypothesis correction, scGeneHE estimates non-zero *cis*-heritability for *KMT2C* exclusively in CD14+ monocytes, supported by a differential gene expression analysis following a weight loss regime; it may be the case that the same *cis*-genetic variants that regulate *KMT2C* also regulate metabolic traits like body mass index (BMI)^88^. The *PAFAH1B1* gene exhibits similar specificity but has not previously been implicated in monocyte regulation.

We also examined the variation of scGeneHE z-scores and standard errors across the myeloid lineage (**Supplementary Figure 9**). Standard errors do not necessarily decrease as we increase the resolution of our cell type clustering. Considering standard errors alone can be misleading, as a gene with a *cis*-heritability estimate of zero (or close to zero) will likely produce a similarly small standard error, because the bootstrapping procedure will tend to consistently estimate near-zero *cis*-heritabilities. As expected, for genes with larger *cis*-heritability estimates, scGeneHE tends to be more confident about these estimates (as opposed to producing overestimates for null genes), reflected by the concordance of z-scores and *cis*-heritability estimates. Overall, we developed a multiscale framework to consider a range of cellular resolutions, which underscores the utility of scGeneHE in uncovering how *cis*-genetic regulation of gene expression operates in both generalizable and cell-type-specific ways.

### Cis-heritability estimation can reveal local genetic architecture obscured by cis-eQTL analysis

We next aimed to compare the information gleaned from *cis*-eQTL association analysis versus *cis*-heritability estimation. *cis*-eQTL analysis complements heritability estimation by identifying discrete SNP-gene expression pairs and estimating a marginal effect size, standard error, and thus p-value. However, only considering these marginal *cis*-eQTL associations can inflate the perceived total genetic effect due to linkage disequilibrium, as is seen for genome-wide association studies^4^. As previously described by the schema in **Figure 1A**, we set out to exemplify the complementary information that is provided by *cis*-heritability estimation. While some previous studies and consortia, including GTEx and OneK1K, define eGenes as those with at least one statistically significant *cis*-eQTL, set-based methods have been widely used to identify even more eGenes, including FastQTL, QTLTools, TensorQTL, APEX, and SAIGE-QTL, as set-based tests can boost power in the absence of a singular strong *cis*-eQTL by aggregating the marginal effects of all *cis*-SNPs^33,17,18,27,89,90,91,92,49^. One such method is the Aggregated Cauchy Association Test (ACAT); ACAT-V calculates gene-level p-values from *cis*-SNP eQTL p-values using Cauchy combinations^93^, specifically used by APEX and SAIGE-QTL. ACAT-V is most powerful under the scenarios of sparse genetic effects (which may be true for gene expression traits) and weak linkage disequilibrium^92^. Therefore, we hypothesized that eGene detection based on set-based tests would not necessarily correspond to genes with significantly nonzero *cis*-heritability estimates, as the set-based tests may be susceptible to the effects of linkage disequilibrium which result in smaller marginal *cis*-eQTL p-values on average. Therefore, *cis*-heritability estimation should provide orthogonal insights regarding the magnitude of the causal genetic component of gene expression.

To this end, we applied the scalable and robust single-cell eQTL association method SAIGE-QTL to the 166 genes considered above in **Figure 4**, e.g., those that are near loci implicated by RA GWAS and that are expressed in all 11 cell types considered in our previous analyses. We obtained *cis*-eQTL summary statistics for each gene and to identify those that were statistically significant, we performed multiple hypothesis correction across all genes and all cell types and employed a 5% FDR significance threshold. We obtained gene-level p-values using the ACAT-V test on nominal *cis*-eQTL p-values, which are inversely weighted by allele frequencies to emphasize contributions from less frequent common alleles. Gene-level p-values are similarly assessed to identify detected eGenes at a 5% FDR significance threshold, correcting for the number of cell types tested for a given gene.

While scGeneHE and SAIGE-QTL identify eGenes via different metrics, we first assessed the concordance of scGeneHE *cis*-heritability p-values with SAIGE-QTL ACAT-V gene-level p-values. Overall, the two methods tend to differ, with scGeneHE indicating that only a small subset of SAIGE-QTL eGenes are actually *cis*-heritable (**Figure 6A, Supplementary Table 9**). Additionally, SAIGE-QTL gene-level p-values tended to be lower on average than scGeneHE *cis*-heritability p-values. We hypothesized that these differences were due to key differences in genetic architecture at these gene loci, such as the number of statistically significant *cis*-eQTLs and the amount of LD across *cis*-SNPs. As demonstrated by previous work^4^, the expectation of the marginal significance of a particular variant (chi-square statistic) is linearly proportional to the total amount of linkage disequilibrium the variant is in with nearby tagging SNPs. While we did not find a strong correlation between either set of gene-level p-values with average linkage disequilibrium in the locus (**Supplementary Figure 10**), we did find that there was a strong correlation between the number of statistically significant *cis*-eQTLs identified by SAIGE-QTL and the strength of the ACAT-V gene-level p-value (R^2^ = 0.41, p-value < 7.4e-10) (**Supplementary Figure 11**). However, as expected, we found that there was not a substantial correlation between the number of significant *cis*-eQTLs identified by SAIGE-QTL and the strength of the scGeneHE heritability p-values (**Figure 6B**, R^2^ = 0.1, **Supplementary Table 9**). Overall, this reinforces our hypothesis that the number of statistically significant *cis*-eQTLs is not a meaningful indicator of *cis*-heritability, which is more strongly influenced by the true unobserved effect sizes of causal eQTLs rather than marginal association statistics.

As depicted in **Figure 1A**, we identified two regimes of possible *cis*-genetic architecture: one in which there is a large number of significant *cis*-eQTLs but low *cis*-heritability and a second in which there are few significant *cis*-eQTL but large *cis*-heritability. A third more intuitive regime that was not depicted is where the number of significant *cis*-eQTLs and the *cis*-heritability of gene expression are both high or are both low. We next explore local genetic architectures in each of these regimes.

First, in the regime of many significant *cis*-eQTLs yet low *cis*-heritability, we exemplify the *PSMB8* gene in intermediate B cells (**Figure 6C**, first row) where there are 5,671 detected significant *cis*-eQTLs (ACAT-V gene p-value < 8.9e-26) but the scGeneHE *cis*-heritability is modest (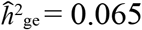, p-value < 2.77e-11). In this case, there may be a small number of causal variants or many causal variants with small effect sizes, but the perceived effect when considering marginal associations is larger due to the extensive LD in this locus.

**Figure 6.**
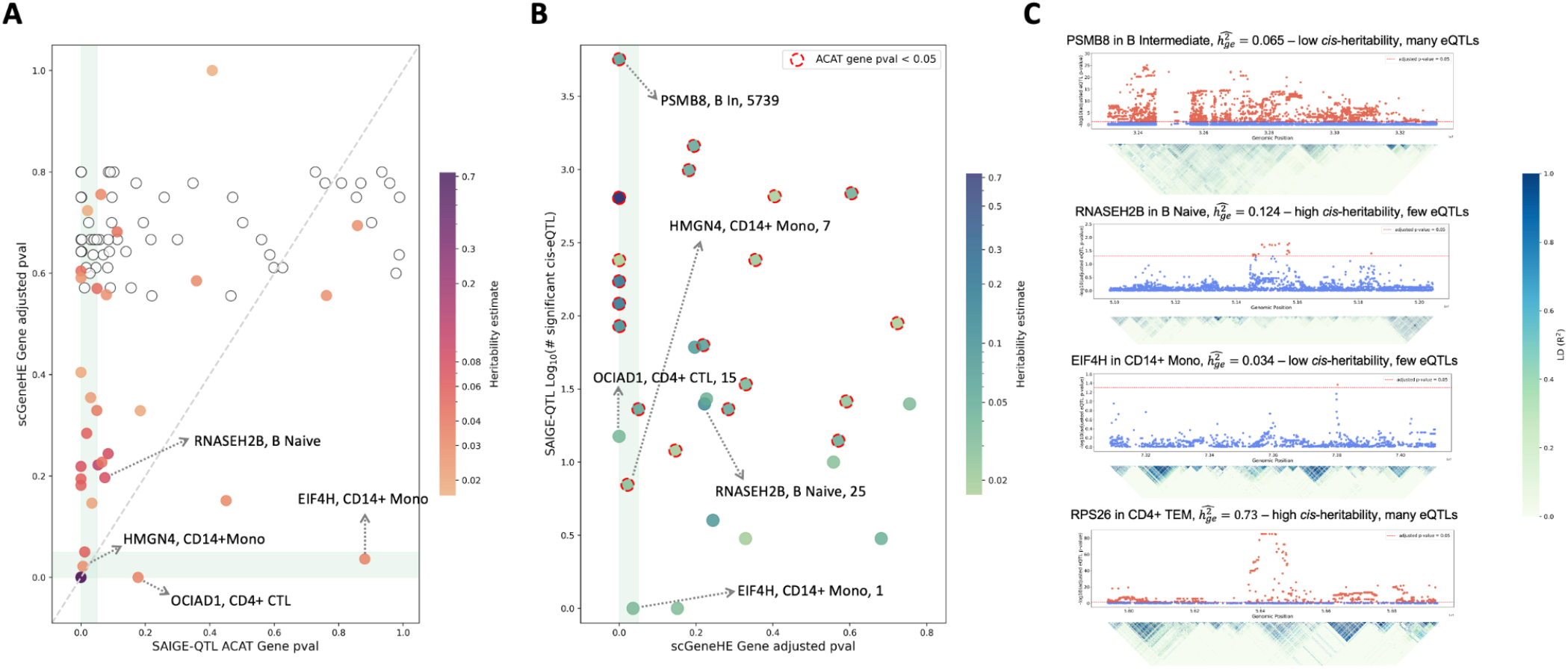
Comparison of gene-level statistics between scGeneHE and SAIGE-QTL. (**a**) FDR-corrected p-values for *cis*-heritability from scGeneHE (y-axis) versus FDR-corrected gene-level p-values calculated by ACAT-V in SAIGE-QTL (x-axis). Vertical and horizontal light green bands indicate FDR < 5% for SAIGE-QTL and scGeneHE, respectively. Data is shown for genes exemplified in **Figure 4**. Color of dots indicates the *cis*-heritability estimate from scGeneHE. White dots with black outline indicate *cis*-heritability estimates of 0. (**b**) Log-scaled number of statistically significant *cis*-eQTLs detected by SAIGE-QTL (y-axis) versus the FDR-corrected *cis*-heritability p-values from scGeneHE (x-axis). Color of dots indicates the *cis*-heritability estimate from scGeneHE. Vertical light green band indicates genes with significantly non-zero estimated *cis*-heritability at 5% FDR. Dotted red outline indicates that the ACAT-V gene-level p-value is significant at 5% FDR. Exemplar genes are annotated with their corresponding cell type and number of detected *cis*-eQTLs. (**c**) LocusZoom plots for four exemplary genes demonstrating that the *cis*-heritability of gene expression is not necessarily reflective of the number of detected *cis*-eQTLs in a gene locus. Numerical results are reported in **Supplementary Table 9**.

Second, in the regime of few significant *cis*-eQTLs and relatively larger *cis*-heritability, we exemplify the *RNASEH2B* gene in naive B cells, which is nominally *cis*-heritable before FDR correction (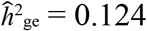 p-value = 0.22, 25 significant *cis*-eQTLs, ACAT gene p-value = 0.053) (**Figure 6C**, second row). In this case, there may be a small number of causal variants with larger effect sizes, although there may be fewer tagging variants to bolster marginal significance, as supported by relatively weaker LD patterns. This regime is particularly helpful in demonstrating the advantage of using scGeneHE to estimate *cis*-heritability and to complement the information gleaned from *cis*-eQTL analysis.

Lastly, we explore the straightforward regime where the *cis*-heritability is concordant with the number of significant *cis*-eQTLs. For instance, *EIF4H* has low *cis*-heritability in CD14+ monocytes (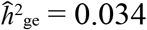, adjusted p-value < 3.6e-2, 1 significant *cis*-eQTL, ACAT gene p-value = 0.88) (**Figure 6C**, third row). Notably, scGeneHE demonstrates the ability to detect *cis*-heritability for a gene that was not considered an eGene by the ACAT-V test, suggesting that it may be a more powerful strategy to aggregate the effects of *cis*-SNPs, especially when these effects are smaller. Conversely, the *RPS26* gene has a large estimated *cis*-heritability in CD4+ T effector memory (TEM) cells (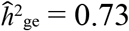, p-value < 1e-300) while also possessing a large number of significant *cis*-eQTLs (*n* = 671) and highly significant ACAT-V p-value (p < 8.6e-121) (**Figure 6C**, fourth row). While extensive LD in the locus may have helped bolster marginal *cis*-eQTL significance, scGeneHE agrees that there are likely several large independent *cis*-eQTL effects in this locus.

Overall, this analysis highlighted a few exemplar genes where loci with more extensive LD tended to more significant *cis*-eQTLs, but this did not always guarantee a larger *cis*-heritability estimate. Therefore, to gauge the magnitude of the true *cis*-genetic component of gene regulation, scGeneHE is likely more informative than marginal *cis*-eQTL associations or set-based gene-level p-values. Notably, in the scenario where there are only a few *cis*-eQTLs detected with weak significance, scGeneHE may still be sufficiently powered to detect a large *cis*-genetic component of heritability. This is particularly advantageous for downstream analyses involving the integration of gene-level information with polygenic disease and complex trait GWAS data, such as transcriptome-wide association studies (TWAS), which routinely overlook and exclude such genes.

## Discussion

In this study, we introduced scGeneHE, a novel Poisson mixed-effects model designed to estimate the *cis*-genetic component of gene expression heritability using scRNA-seq data. This method uniquely addresses the challenges posed by the high sparsity, non-independence of measurements, and non-normal distributions of read counts from scRNA-seq experiments, providing a robust framework for understanding the genetic regulation of gene expression at unprecedented resolution. scGeneHE addresses a critical gap in the field of single-cell genomics: scGeneHE is the first method to explicitly model intra-individual cell dependencies for heritability estimation, by incorporating a cell-cell relatedness matrix (CRM) alongside a genetic relationship matrix (GRM) into a Poisson mixed-effects framework. This approach not only improves the accuracy of heritability estimation but also enables the identification of cell-type-specific genetic regulation with well calibrated type I error, which greatly contributes to understanding the molecular mechanisms that disease-associated variants may be involved in. We note that other methods have been recently developed to estimate local *cis*-heritability of genome-wide complex traits^6^ and *cis*-heritability of gene expression from pseudobulk scRNA-seq data^11^, but neither leverages multiple molecular measurements per donor.

Through extensive simulations, we evaluated the bias for causal and null *cis*-heritability estimation, power, and type I error rate for eGene detection using scGeneHE or the pseudobulk approach. Specifically, we evaluated model performance across varying levels of true *cis*-heritability, numbers of causal *cis*-eQTLs, donor sample size, cells per donor, sparsity, and cellular heterogeneity, which are the most defining features of scRNA-seq experiments. Most notably, scGeneHE maintained controlled false positive rates on eGene detection across an array of parameters. We found that scGeneHE performs as expected when there is variation in shared cellular environments (reflecting different levels of clustering of real cellular profiles). In contrast, the pseudobulk approach, which treats cells as independent measurements, more frequently overestimates heritability and consistently exhibits staggeringly high false positive rates. We find that the tradeoff of scGeneHE having lower eGene detection power than the pseudobulk approach is justified by its reliable type I error calibration.

We then applied scGeneHE to 11 immune cell types and 366 total genes in the OneK1K cohort, identifying 82 gene-cell-type pairs with significant cell-type-specific genetic regulatory components. scGeneHE identifies eGenes in biologically plausible cell types with established roles in immune system regulation, particularly illuminating how genes like *PKIG* and *ZFP36L1* are genetically regulated within specific immune cell subtypes such as naive B cells and CD16+ monocytes, respectively. For these and many other genes, the pseudobulk approach either implicated many cell types without specificity or implicated a less biologically plausible cell type. scGeneHE also captured new eGenes that were not detected by the pseudobulk approach, including *RNASEH2B* in naive B cells and *ILF3* in CD8+ naive T cells. As opposed to univariate *cis*-eQTL association testing, scGeneHE identifies eGenes by jointly modeling contributions from all *cis*-SNPs. Our method complements marginal *cis*-eQTL association analysis and set-based tests which are commonly used to identify eGenes, as these tests can be misleading because marginal associations are so strongly influenced by local LD. For instance, scGeneHE estimated that *PSMB8* has low *cis*-heritability in intermediate B cells although this gene harbors 5,739 statistically significant *cis*-eQTLs and conversely, scGeneHE found that *RNASEH2B* has relatively high *cis*-heritability in naive B cells although the locus harbors only 25 *cis*-eQTLs across two LD-independent peaks.

We note several limitations of our work. First, the method assumes a simple identity matrix for the CRM, which implies that all cells from the same individual share a perfectly correlated environment. While this assumption simplifies the model and makes it computationally tractable, it may not fully capture the complexity of cellular heterogeneity within individuals due to differences in spatial cellular environments and cell-cell signaling. However, from the perspective of model design, a more complex CRM could lead to a random slope model, which would complicate the estimation process and potentially introduce instability. Future work could explore alternative approaches to model cell-cell relatedness more accurately, such as incorporating cell-type-specific covariance structures.

Second, the current application of scGeneHE is limited by the availability of real-world data with sufficient relatedness between individuals. Contrary to the standard desire for unrelated individuals in genetic association analysis, minor relatedness is actually useful in our framework to decompose phenotypic variation into inter- and intra-individual variation. In the OneK1K cohort, the average relatedness between individuals is relatively low (approximately 20% correlation across genotypes), resulting in a relatively flat likelihood function where the optimizer struggles to find the global maximum. This results in reduced power to detect *cis*-genetic effects, especially affecting genes with small *cis*-heritability. Future cohort recruitment with greater relatedness among individuals could improve the performance and utility of scGeneHE.

Third, scGeneHE assumes an ideal Poisson distribution for gene expression counts. While the Poisson distribution is a reasonable approximation for scRNA-seq data, real-world data often exhibit overdispersion or zero-inflation, which could be better captured by negative binomial or zero-inflated models^50^. Deviations from the Poisson assumption could affect the fitting of scGeneHE and lead to biased heritability estimates. Although in the model design we fix the dispersion parameter to be 1, relaxing it does not provide much freedom to fit a distribution of read counts that is not perfectly Poisson. Future extensions of scGeneHE could incorporate more flexible count distributions to better accommodate the complexities of scRNA-seq data.

Fourth, we employ an empirical bootstrap to determine *cis*-heritability standard errors, for lack of a theoretical standard deviation. Due to the complexity of the Poisson mixed-effects model and the use of optimization and approximation techniques in solving the restricted maximum likelihood (REML) equations, deriving a theoretical standard deviation is statistically challenging. As bootstrapping requires repeatedly refitting the model to resampled data, this can significantly increase the runtime of the analysis, making it less practical for larger datasets.

Finally, while scGeneHE demonstrates robust performance in simulations and real data analyses, its computational efficiency remains a challenge, particularly for large-scale datasets with millions of cells and thousands of genes. We estimate that the time to run scGeneHE scales linearly with the number of cells in a dataset. Although the method leverages efficient algorithms for mixed-effects models, even further optimization will be necessary to make it scalable to conduct bootstrapping for thousands of donors and tested genes.

In summary, scGeneHE represents a significant advancement in the estimation of gene expression heritability from scRNA-seq data. By addressing the limitations of pseudobulk approaches and providing a framework for cell-type-specific heritability estimation, scGeneHE opens new avenues for understanding the *cis*-genetic regulation of gene expression and how it may contribute to the mechanisms of complex traits and polygenic diseases. While we explore several limitations of our method, its ability to detect cell-type-specific genetic components with controlled false positive rates makes it a valuable tool for interpreting single-cell genomics data.

## Methods

### scGeneHE Model

scGeneHE uses a Poisson linear mixed model to address the sparsity and repeat measurements in single cell RNA-seq gene expression data. Our model formulation refers to a total of n individuals and M cells across individuals, where each individual *i* has m_i_ cells, such that 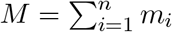. We model the sequencing read count of a specific gene, *y*_ij_, using a Poisson distribution, such that *y*_*ij*_ *~ Pois*(*µ*_*ij*_), where j indexes cells from the i-th individual. To ensure the Poisson parameter value µ_ij_ is always positive, we use a linker function *µ*_*ij*_ = *g*(*µ*_*ij*_) = log(*µ*_*ij*_), as previously employed^94^. Now, the additive, linear relationship between the genetic and environmental components of gene expression exists in the logarithmic transformed space of our Poisson parameter *µ*_*ij*_. We let 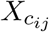 represent cell-level covariates such as gene expression PCs, mitochondrial percentage and total normalized UMI count, where *i* and *j* index the covariate vector for individual *i* and cell *j*. We let 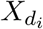 represent individual-level covariates such as genotype PCs, sex, age and pool number, where *i* indexes the covariate vector for individual *i*. The linear relationship given by our linker function is thus the following: 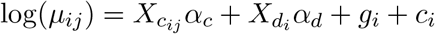 where the first two covariate terms are fixed effects. The random effect g_**d**_ is a vector of length n where each element is g_i_; g_**d**_ is assumed to follow a multivariate normal distribution g_**d**_ *~ N*(0, τ_1_A), where τ_1_ is the true unobserved variance parameter of the genetic component and A is the observed n *×* n individual genetic relationship matrix. The random effect c_**d**_ is a vector of length n where each element is *c*_*i*_; c_**d**_ is assumed to follow a multivariate normal distribution c_**d**_ *~ N*(0, τ_2_I_**n**_), where τ_2_ is the true unobserved variance parameter of the cell-cell relatedness component and I_**n**_ is the *n × n* identity matrix representing intra-individual cellular relatedness. Since we assume g_**d**_ and c_**d**_ are independent, we may condense both random effect terms into one general term ***θ***_d_ where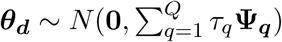. In our specific case, Q = 2, Ψ_**1**_ = A, and Ψ_**2**_ = I_**n**_.

The read counts *y*_*ij*_ are assumed to be independent, conditional on the random effects g_i_ and c_i_. They follow a Poisson distribution with mean E(*y*_*ij*_ | θ_i_) = µ_ij_ and variance ϕ Var(*y*_*ij*_ | θ_i_) = ϕ *var*(µ_ij_) = µ_ij_, where is the dispersion parameter which is set to 1 by default.

Next, we describe the matrix that designates which cells belong to which individuals. Let **Z** be an *M × n* design matrix. Each row in **Z** contains a singular 1, to indicate the individual to which the cell belongs, while all other elements are set to 0. Specifically, for each *i* ∈ {1, 2,…, n} and *j* ∈ {1, 2,…, *M*}, the row *Z*_*i*,*j*_ satisfies:

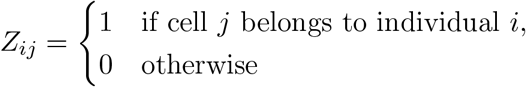

**Z** could also be visualized as a matrix of n virtually concatenated matrices **Z**_**i**_ of size *m*_i_ *× n*, where each

**Z**_**i**_ represents cell membership of the *i*-th individual. Then the Poisson mixed model can be written as

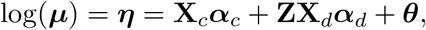

where log ***µ*** is a vector of length *M*, X_c_ is a matrix of size *M ×* (*p*_c_ + 1) containing cell-level covariates with intercept, X_d_ is a matrix of size M *× p*_d_ containing individual-level covariates, and the random effects for all cells **θ** = **Z** θ_d_ is a vector with length *M*. Specifically, the two random effects g and b of length *M* will have observed variance-covariance matrices **ZAZ**^T^ representing the expanded cell-level genetic relatedness matrix and **ZI**_**n**_**Z**^T^ representing the cell-cell relatedness matrix, which indicates that cells from different individuals are unrelated.

Based on previous work^57^, when **ZAZ**^T^ is standardized to have *tr*(**ZAZ**^*T*^)/*M* = 1, heritability will be constrained to the interval [0, 1] and will now be comparable to linear space heritability, where the tr(·) denotes the trace of a matrix. We then define heritability as

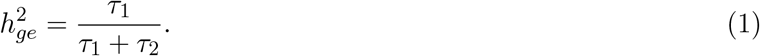

### Penalized quasi-likelihood

We adapt the same process of fitting a penalized quasi-likelihood from SAIGE-QTL^49^. To estimate (α, *ϕ*, τ) where *α* = (*α*_c_, *α*_d_) and τ = (τ_1_, τ_2_), the integrated quasi-likelihood function is:

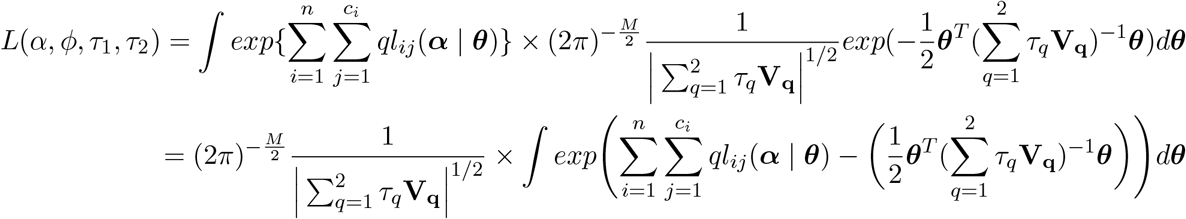

where V_**q**_ = Z Ψ_*q*_Z^*T*^,

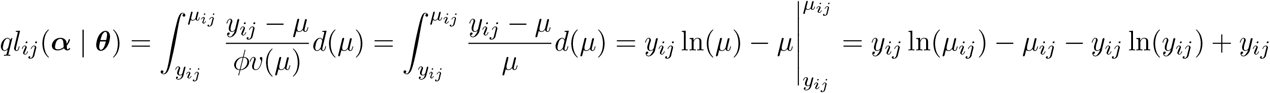

is the quasi-likelihood for the j-th cell from the i-th individual given the random effects θ.

If we let 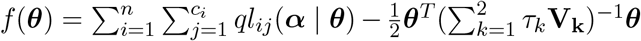, we can then approximate the integral using the Laplace approximation,

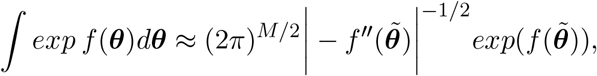

where 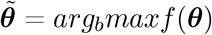 is the solution to f ^*′*^ (***θ***) = 0. Therefore, the log-integrated quasi-likelihood function can be approximated by

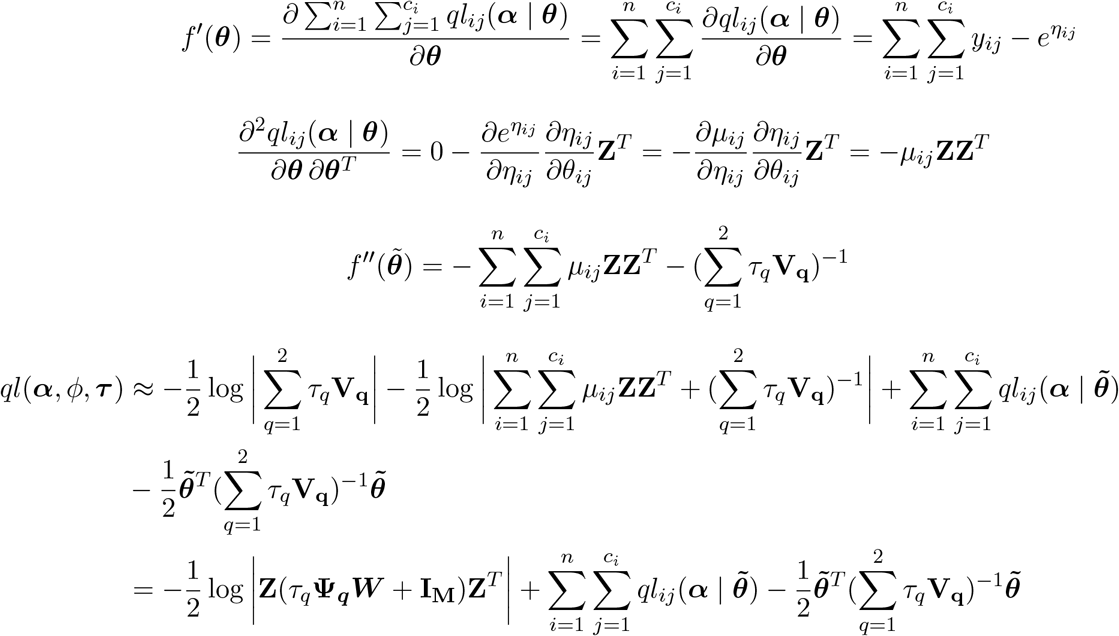

where **W** = *diag*(*µ*_*ij*_).

### Estimation of fixed and random effects using AI-REML

To obtain the estimates of the fixed effect coefficients and random effects given (ϕ, **τ**) and 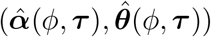 that jointly maximize ql(**α**, *ϕ*, **τ**), we take the derivative of ql(**α**, *ϕ*, **τ**) with respect to **α** and **θ** and solve for the parameters that make the derivatives zero.

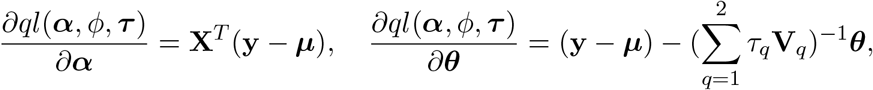

where **X** = [**X**_c_, **ZX**_d_]. Let **Y** be a working vector **Y** = **η** + **W**^−1^(**y** *−* ***µ***) where **W** = *diag*(***µ***_ij_), **η** = (*η*_1_, …, *η*_*M*_), and ***µ*** = (*µ*_1_, …, *µ*_M_). Then we have **y** *−* ***µ*** = **W**(**Y** *−* η) = **W**(**Y** *−* **X**_α_ *–* θ). Then, the solution of derivatives may be written as:

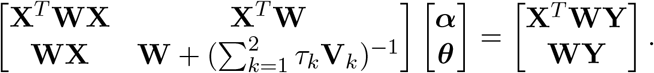

If we let variance matrix 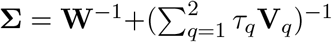 and projection matrix **P** = **∑**^−1^−**∑**^−1^**X**(**X**^*T*^ **∑**X)^−1^X^*T*^ **∑**^−1^, we have

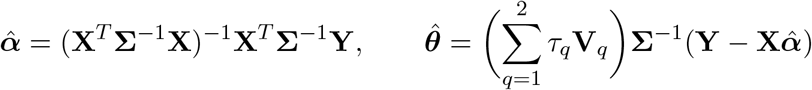

Given the estimated values of 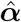 and 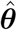,

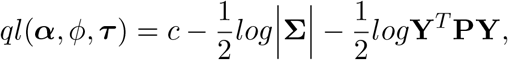

then the restricted maximum likelihood (REML) is

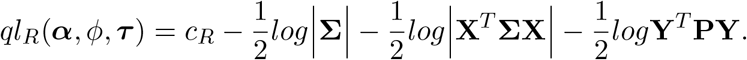

Thus, the derivative of the REML likelihood function with respect to τ is

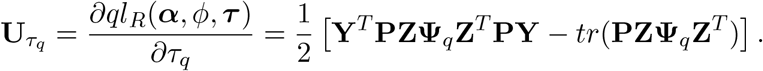

The corresponding observed information function and expected information matrices are

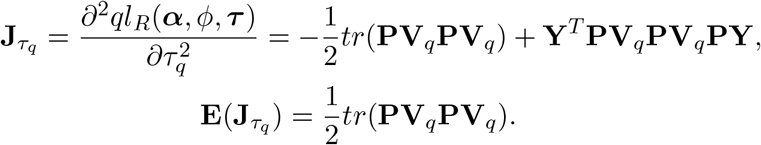

The average information matrix is

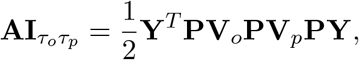

where AI is a 2 *×* 2 matrix with (o, p)-th element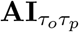.

### Hypothesis testing and standard error estimation using bootstrap

We iteratively fit the scGeneHE model to obtain mean estimates of the variance parameters and point estimate of heritability 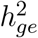 via equation (1),

1. Fit a Poisson linear model with **τ** = 0 to get initial estimates 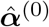 and working outcome vector **Y**^0^
2. At the i-th step, update 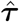 using 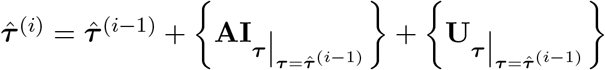
3. Update 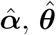 using **Y** and 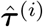
4. Update **Y** using 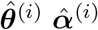, and 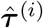
5. Repeat steps 2–4 until 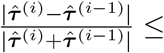 *tolerance*.

To obtain standard errors of our point estimates, we conduct a non-parametric bootstrap by sampling *M* cells with replacement 100 times. For each bootstrap, we estimate the variance parameters **τ** ^*\**^ and heritability 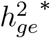 with the same process above. Bootstrap heritability is assumed to follow 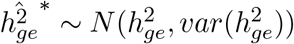 and the bootstrap standard deviation 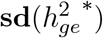 is assumed to be an approximation of the empirical standard deviation of heritability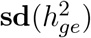.

To determine a significantly heritable gene, we use a one-sided t-test to test the null hypothesis 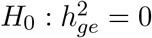 against the alternative hypothesis 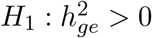 given significance level *α*, using the point estimate 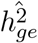 and bootstrap standard error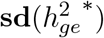.

### Simulations

We simulated hundreds of iterations of various genetic architectures using real SNP genotypes to evaluate the power, bias (causal and null), and calibration of scGeneHE. We focus on a 1 Megabase (Mb) region of chromosome 1 using the genotypes of n randomly selected European individuals from phase 3 of the 1000 Genomes Project. For each individual, we simulate m cells and assume that cells of one cell type within each person share the same genetic component (predefined *cis*-heritability of gene expression 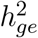) but variable environmental noise sampled from a distribution of 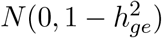 (see below for more details). For each individual, we sample *n*_*causal*_ eQTL effect sizes from a normal distribution of mean zero and variance 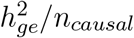, where all non-causal SNPs are defined to have effect sizes of zero. For each independent simulation (typically 100 simulations in total per experiment), we randomly sample new causal eQTL effect sizes for a gene from the distribution described. Then for each simulation, we generate the individual-level *cis*-genetic component of gene expression as the dot product of the standardized genotype matrix of *cis*-SNPs and the sampled eQTL effect size vector. All m cells of a given cell type from a given donor share the same *cis*-genetic component of gene expression.

Then, we introduce the index of cellular heterogeneity *K* to sample cell-level environmental noise, to reflect more realistic scenarios of the existence of cellular subtypes within a given cell type and fluctuations of environmental noise based on spatial position or cell-cell signaling. For example, if *K* = 1, all cells of a given cell type from a given donor have the same sampled environmental noise added to the *cis*-genetic component of gene expression. We first randomly split the m cells into *K* groups such that there are *m/K* cells per group (rounded up to the nearest integer). Each group of cells is assigned the same environmental noise ϵ by sampling a single value from a normal distribution of mean zero and variance equal to 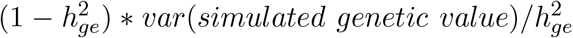. By taking the exponential (inverse of linker function of Poisson mixed model) of the sum of genetic value and environment noise, we generate non-negative single-cell gene expression. Then, we randomly substitute r% of cells for each person to be zero, to impose sparsity representing the rates of technical zeros (or drop-out) in single-cell RNA-seq data^53^.

For each independent simulation of single cell gene expression with a non-zero *cis*-genetic component, we also simulate corresponding single cell gene expression without a *cis*-genetic component (null analysis) and, separately, pseudobulk gene expression values. Null single cell gene expression data is generated by randomly permuting cell-donor assignment of the single cell expression matrix with a non-zero *cis*-genetic component; this disrupts the association between gene expression and genotype. We note that when the sparsity rate is high (e.g. > 80%) and thus there are many more zeros than non-zero values in the single cell gene expression matrix, this permutation strategy may not be effective at disrupting associations between gene expression and genotype, as the assignment of zeros after permutation may substantially overlap with the original assignment of zeros. Under this specific scenario, we have observed slightly inflated null bias on the estimation of 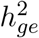; however, the type I error remains well-calibrated. Next, we simulate pseudobulk gene expression for the scenario of true *cis*-heritable genes, as well as null genes, by taking the average or sum of gene expression across cells within each person.

Constructing a genetic relationship matrix (GRM) is critical to modeling inter-individual variation and thus the estimation of heritability. We create an (*m × n*) by (*m × n*) cell-level GRM by duplicating the rows of the canonical individual-level GRM *m* times. To decompose the variance in total gene expression into genetic inter-individual variation and intra-individual variation between cells, we construct a cell-cell relatedness matrix (CRM) as a block diagonal matrix 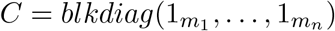 representing the binary assignment of cells to the corresponding donor. We conducted fast principal components analysis (PCA) analysis on genotypes and gene expression values using the R package ‘irlba’ (v2.3.5.1) and use the top 6 genotyping PCs as covariates, as previously done^27^ for the OneK1K scRNA-seq dataset that we re-analyze in this manuscript. To best mimic real data analysis, we also incorporate mitochondrial gene percentage of each cell as a covariate by randomly sampling values from the following distribution: *Unif* (0, 1).

For each simulated gene in our *cis*-heritable and null scenarios, we estimate heritability using scGeneHE and conduct 100 bootstraps across single cell measurements (enforcing equal donor sample size as to not lose power) and obtain empirical standard errors of the *cis*-heritability of gene expression and the variance attributed to cell-cell relatedness. We detect significantly *cis*-heritable genes by conducting a t-test with a null hypothesis where 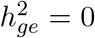 and alternative hypothesis 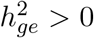 with a nominal significance p-value threshold of 0.05. For both *cis*-heritable and null genes, power and type I error to detect heritability are defined as the percent of simulations where the proportion of gene expression variation attributed to inter-individual variation (e.g., the GRM) is significantly greater than 0.

For each *cis*-heritable and null pseudobulked gene, we standardize gene expression values and estimate *cis*-heritability using the same top 6 genotyping PCs as covariates using GCTA^3^. We use the analytical standard error estimates from GCTA to detect significantly *cis*-heritable genes and calculate power and type I error rates.

We also used ROC curves to assess the relationship between the sensitivity (power) and specificity (one minus the type I error rate) across 5 gene expression heritability values for both single-cell and pseudobulk expression profiles across 1,000 uniformly spaced p-value thresholds.

Through all simulation analyses, we varied 5 parameters to investigate the impact on heritability estimation by changing one while controlling the others one at a time: heritability (0%, 5%, 10%, 25%, 50%), number of people (50, 100, 500), number of cells per person (25, 50, 75, 100), number of causal eQTLs (1, 2, 5), single-cell sparsity (0%, 10%, 40%, 80%, 90%), and shared cellular environment (K = 1, 2, 5, 7).

### Pseudobulking in real data

We generated pseudobulk expression profiles for genes in the OneK1K dataset. We use the transcripts per million (TPM) method to generate normalized scRNA-seq counts that account for gene length and per-cell library size. We first divide the UMI counts for each gene by the length of the gene in Kilobase pairs, as longer genes will tend to have more read counts. Then we scale each cell’s expression profile across genes to sum to 1 million total UMI counts (e.g., TPM). Then we either add up or take the average of TPM counts across cells within each individual to generate individual-level expression counts. We then take the log 2 base transformation of individual-level expression counts plus one, e.g., log_2_(TPM + 1), which serves as the final individual-level pseudobulk expression counts.

### Statistics and Reproducibility

All down-sampling of cells was performed randomly and within each individual. All sampling with replacement used for bootstrapping was performed randomly across cells within each individual. Randomization and blinding were not pertinent to our study.

## Data Availability

GWAS summary statistics are available at https://www.ebi.ac.uk/gwas/efotraits/EFO_0000685. Gene expression and genotype data from the OneK1K cohort are available at the Gene Expression Omnibus (GEO), accession GSE196830. GTEx gene expression and genotype data were acquired from dbGaP accession phs000424.v9.p2.

## Code Availability

scGeneHE software including documentation and tutorial is publicly available at https://github.com/AmariutaLab/scGeneHE [Zenodo DOI https://doi.org/10.5281/zenodo.14920579]. GCTA is available at https://yanglab.westlake.edu.cn/software/gcta/. SAIGE-QTL is available at https://github.com/weizhou0/qtl.

## Acknowledgements

This work was supported by funding from the National Science Foundation (NSF) (Award Number 2336469 awarded to T.A.) and the National Institutes of Health (NIH) (NHGRI R01HG013671 awarded to T.A.). The funders played no role in study design, data collection and analysis, decision to publish or preparation of the manuscript. The Genotype-Tissue Expression (GTEx) Project was supported by the Common Fund of the Office of the Director of the National Institutes of Health, and by NCI, NHGRI, NHLBI, NIDA, NIMH, and NINDS. This work used the Expanse HPC server at the San Diego Supercomputer Center (SDSC) through allocation BIO230210 from the Advanced Cyberinfrastructure Coordination Ecosystem: Services and Support (ACCESS) program, which is supported by National Science Foundation grants 2138259, 2138286, 2138307, 2137603, and 2138296. We are thankful for all participants in the OneK1K study cohort whose data we analyzed in this manuscript.

## Author contributions

Z.X. and T.A. conceived and designed the study. Z.X. conducted simulation analyses and real data analyses. A.R.M. provided support with software environments and reproducibility. L.R. and S.R. provided pre-processed gene expression and genotype data. M.G. provided feedback on the study design. W.Z. helped design model framework and simulation analyses. Z.X. and T.A. wrote the initial draft of the manuscript and all co-authors contributed to the final manuscript.

## Competing Interests

The authors declare no competing interests.

## Notes

### Competing Interest Statement

The authors have declared no competing interest.

