## Supplementary Note for "Estimating the *cis*-heritability of gene expression using single cell expression profiles controls false positive rate of eGene detection"

### *Supplementary Figures*

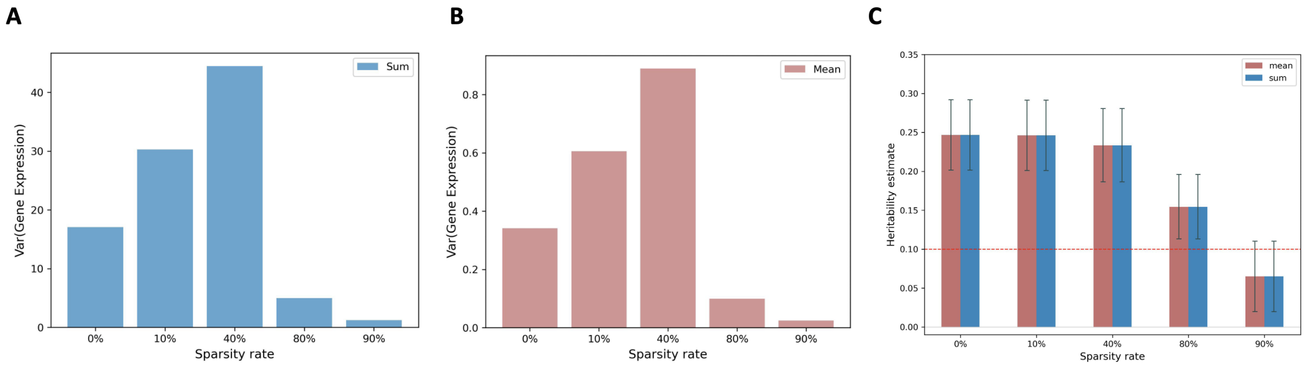


**Supplementary Figure 1: Comparison between averaging and aggregating cell counts within individuals in pseudobulk.** We illustrate the variance of total simulated gene expression when summing (**a**) versus averaging (**b**) cell counts to generate pseudobulk profiles using one set of simulated data: true *cis*-heritability is 0.1, 100 donors in total, 50 cells per donor, 5 causal eQTLs, and varying sparsity rate. The varying sparsity rate results in the same trend in the variance of gene expression; we note that the scale of variance of total simulated gene expression differs between the sum and mean approaches but produces similar proportional changes in variance. (**c**) GCTA generates identical *cis*-heritability estimates for both mean and sum approaches. Black line segments indicate the 95% confidence interval calculated using the standard deviation of the estimates across 100 independent simulations. For consistency, we elect to use the sum approach for the pseudobulk approach in our remaining analyses.

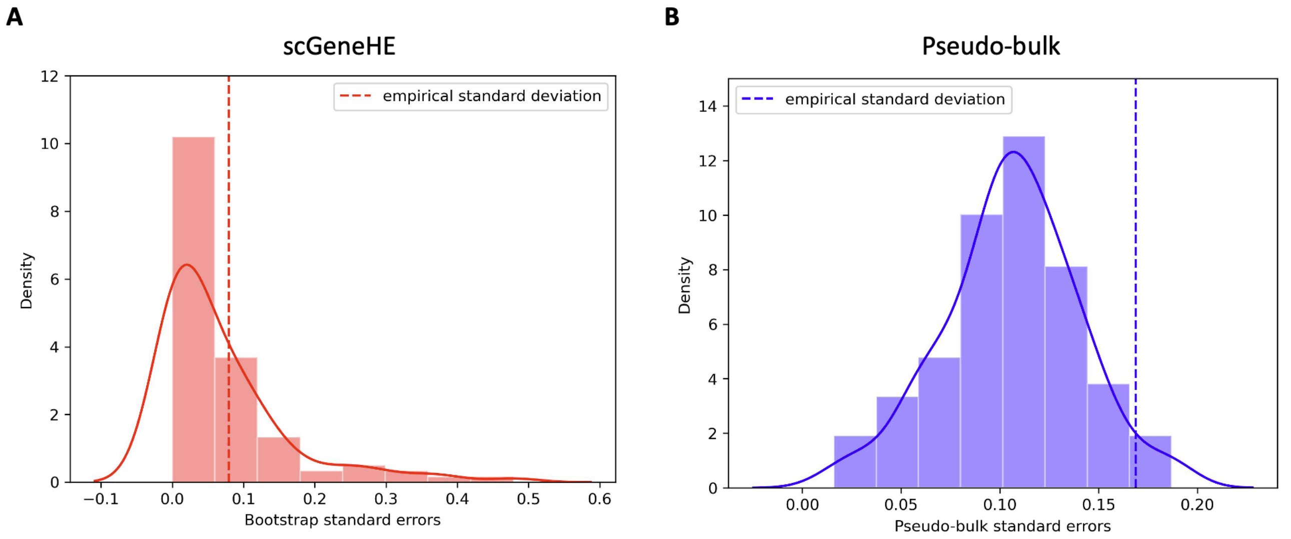


**Supplementary Figure 2: Distribution of estimated standard errors from *cis*-heritability analysis versus distribution of empirical standard deviations of *cis*-heritability estimates across simulations.** (**a**) scGeneHE and (**b**) pseudobulk standard errors were calculated from the same simulations of genes with no true *cis*-genetic components, derived from permutation analysis. The following set of parameters were fixed: 100 donors, 50 cells per donor, $K = 5$, sparsity rate=80%. scGeneHE bootstrap standard errors (**a**) are collected from 100 bootstrap samples of 1 gene. The empirical standard deviation of *cis*-heritability estimates is calculated from 100 independent simulations (red dashed line); because this value falls within the distribution of bootstrap standard errors, we believe our approach to calculating standard errors is well-calibrated. Pseudobulk (**b**) standard errors are analytical estimates generated by GCTA across 100 simulations and the dashed blue line represents the standard deviation of GCTA *cis*-heritability estimates across these same simulations. We note that the GCTA standard errors are seemingly miscalibrated and smaller than expected, which may contribute to the large false positive rates we observed in **Figure 2**.

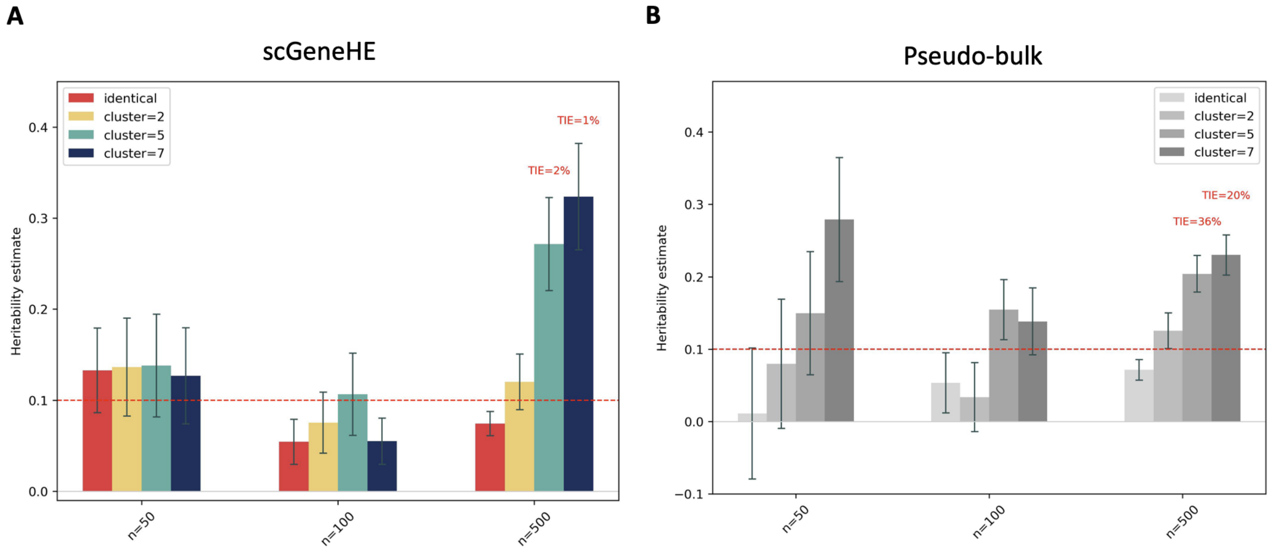


**Supplementary Figure 3:** ***Cis*-heritability estimates as a function of donor sample size.** (**a**) scGeneHE and (**b**) pseudobulk *cis*-heritability estimates for different values of $K$ across varying donor sample size (n, x-axis). Smaller values of $K$ represent more similar environment for cells belonging to the same individual. The following parameters are fixed across settings: 10% true *cis*-heritability (dashed red line), 5 causal eQTLs, 50 cells per donor, and 80% sparsity rate. Black line segments represent the 95% confidence intervals, using the standard deviation across 100 independent simulations. Type I error is annotated for overestimates of heritability by scGeneHE to show that false positive rate is still controlled.

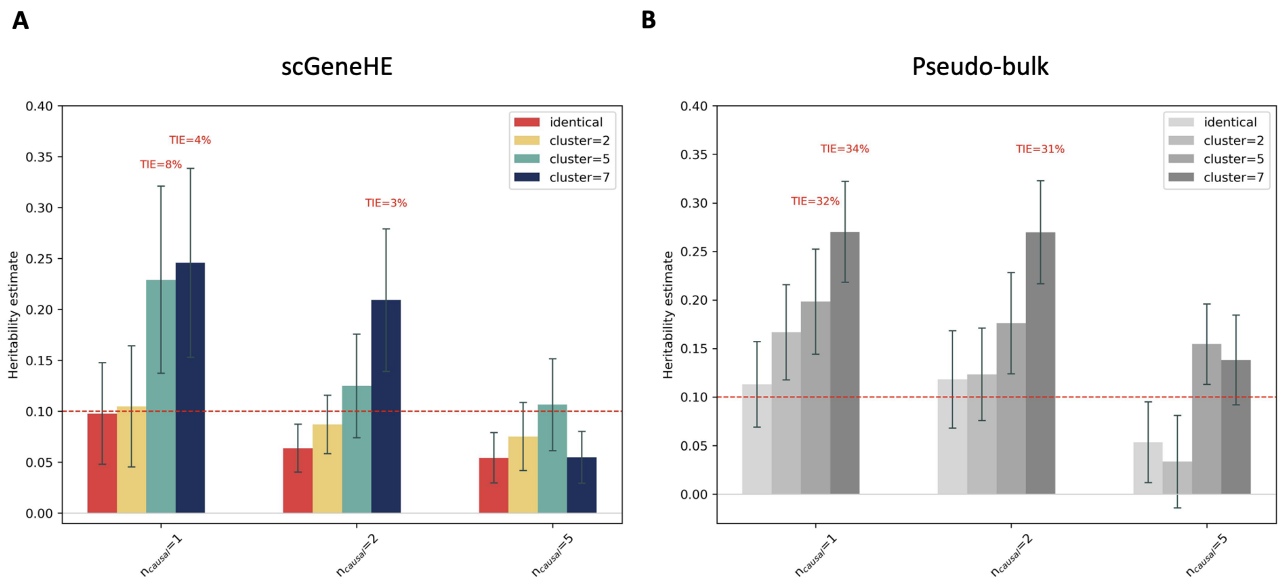


**Supplementary Figure 4: *Cis*-heritability estimates as a function of the number of causal *cis*-eQTLs.** (**a**) scGeneHE and (**b**) pseudobulk *cis*-heritability estimates for different values of $K$ across varying numbers of causal *cis*-eQTLs. The following parameters are fixed across settings: 10% true cis-heritability (dashed red line), 100 donors, 50 cells per donor, and 80% sparsity rate. Black line segments represent the 95% confidence intervals, using the standard deviation across 100 independent simulations. Type I error is annotated for overestimates of heritability by scGeneHE.

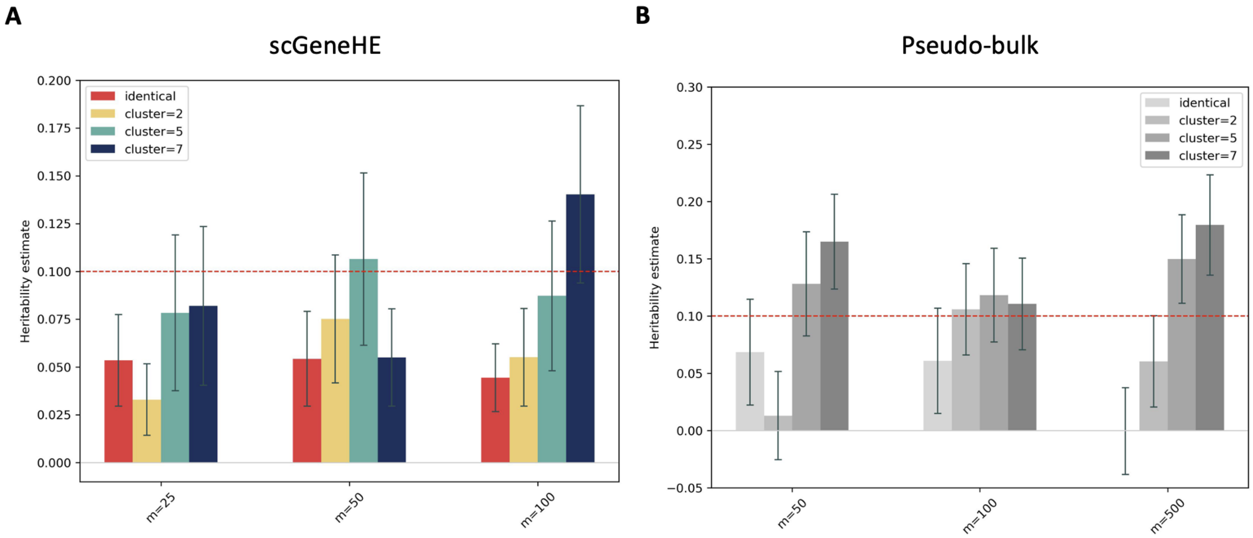


**Supplementary Figure 5:** ***Cis*-heritability estimates as a function of the number of cells per donor.** (**a**) scGeneHE and (**b**) pseudobulk *cis*-heritability estimates for different values of $K$ across various numbers of causal *cis*-eQTLs. The following parameters are fixed across settings: 10% true *cis*-heritability (dashed red line), 100 donors, 5 causal eQTLs per gene, and 80% sparsity rate. Black line segments represent the 95% confidence intervals, using the standard deviation across 100 independent simulations.

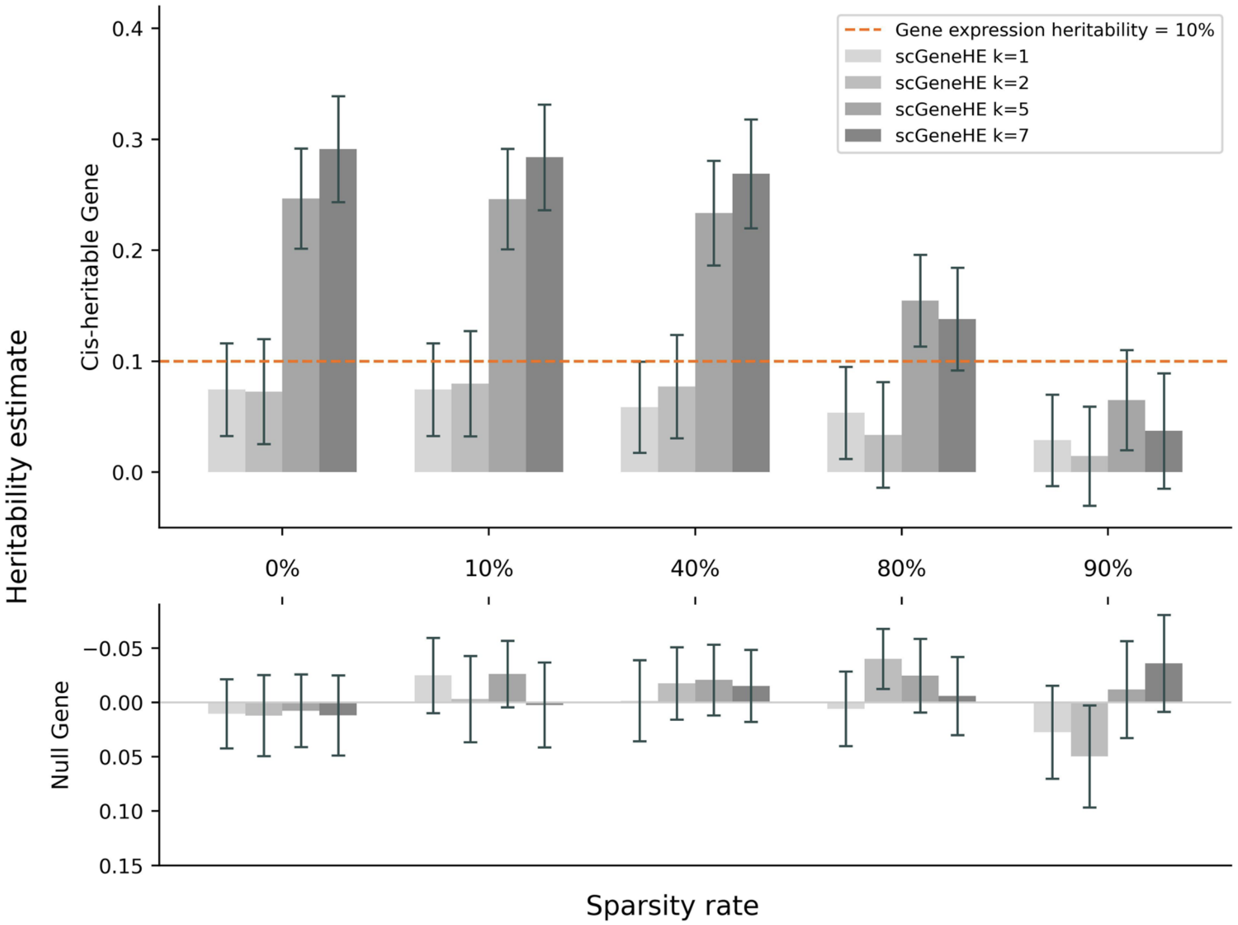


**Supplementary Figure 6: Pseudobulk *cis*-heritability estimates across different sparsity rates and heterogeneity levels.** Pseudobulk *cis*-heritability estimates for different values of $K$ across various levels of sparsity (**top**). Black line segments represent the 95% confidence interval calculated from the standard deviation across 100 independent simulations. Inverted bar graph depicting the *cis*-heritability estimates for genes with no true *cis*-genetic component, derived from permutation analysis (**bottom**).

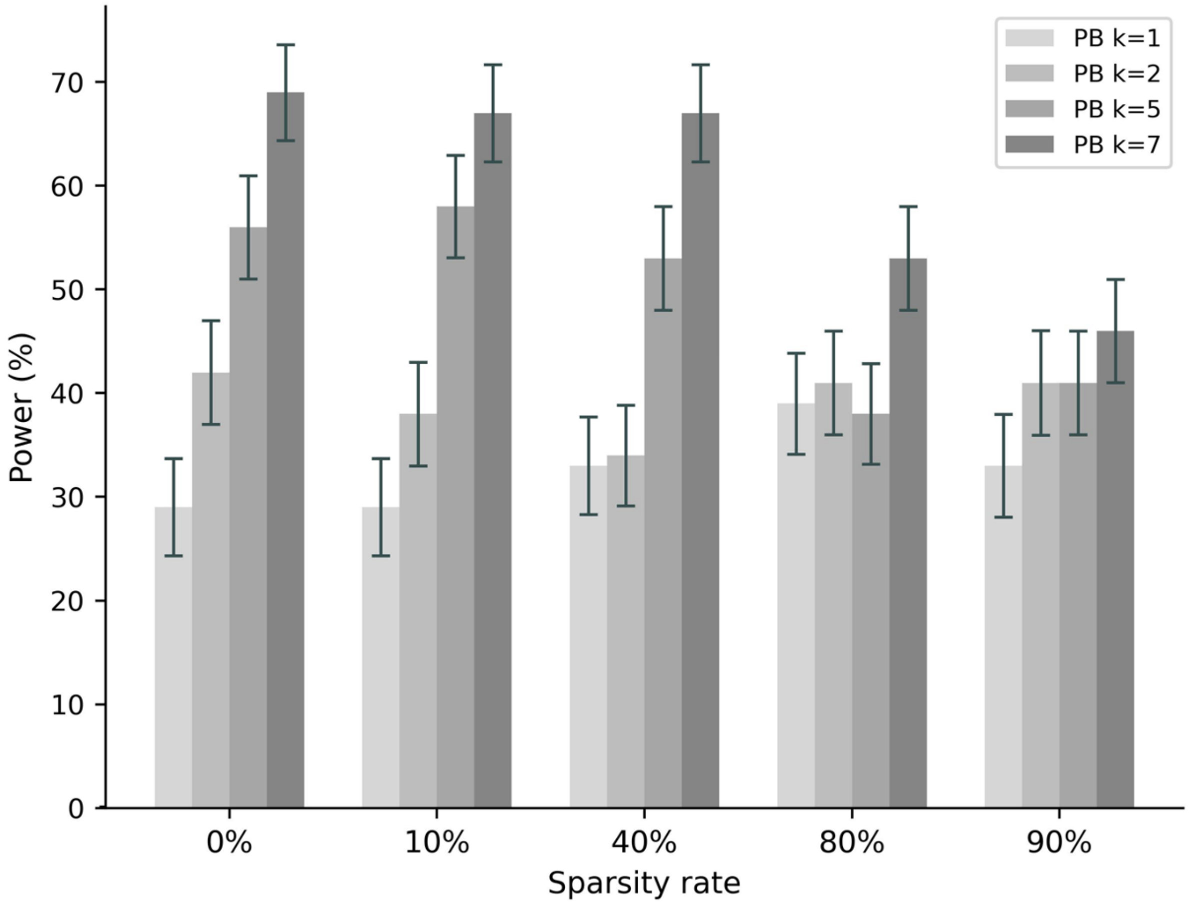


**Supplementary Figure 7: Pseudobulk power across sparsity rates and heterogeneity levels.** Black line segments represent the 95% confidence interval calculated from the standard deviation of power across 100 independent simulations.

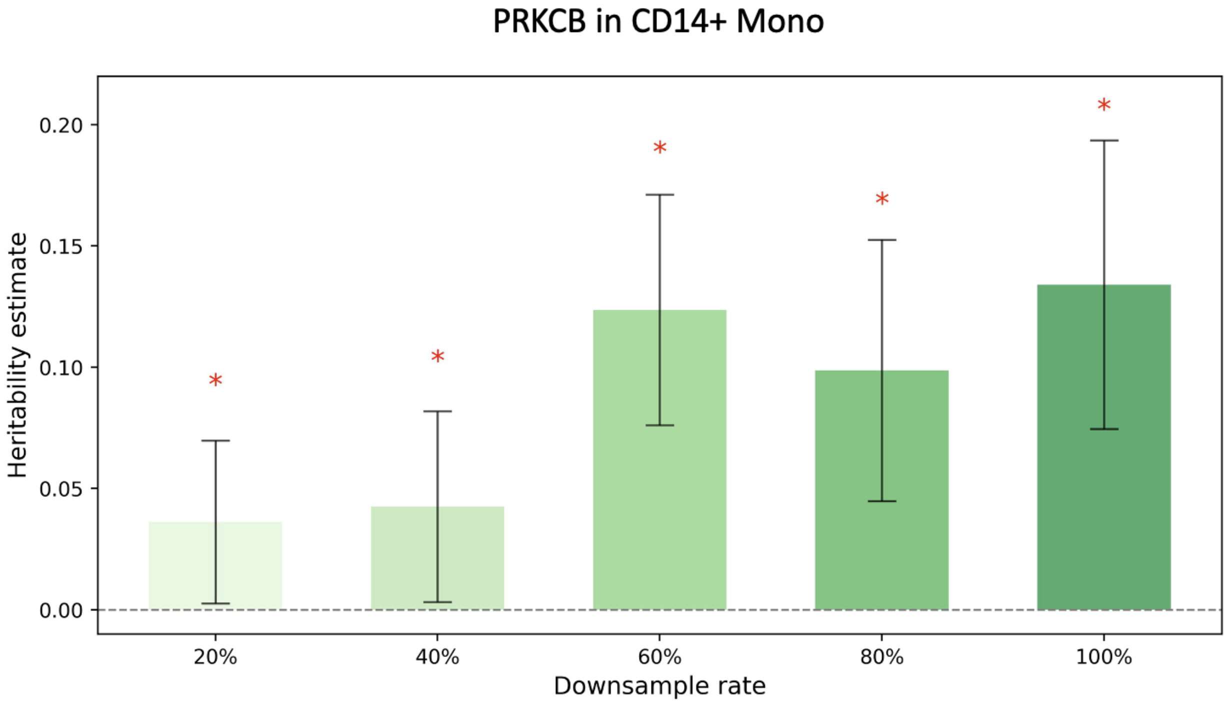


**Supplementary Figure 8: *cis*-heritability estimates of *PRKCB* in down-sampled CD14+ monocytes.** All cells *PRKCB* expressed in CD14+ monocytes were collected to conduct a random downsampling. For each level of downsampling, the *cis*-heritability estimate was significantly non-zero, indicated by a red asterisk at the top of the bar. Bars represent the 95% confidence interval calculated from the bootstrap standard error calculated by scGeneHE.

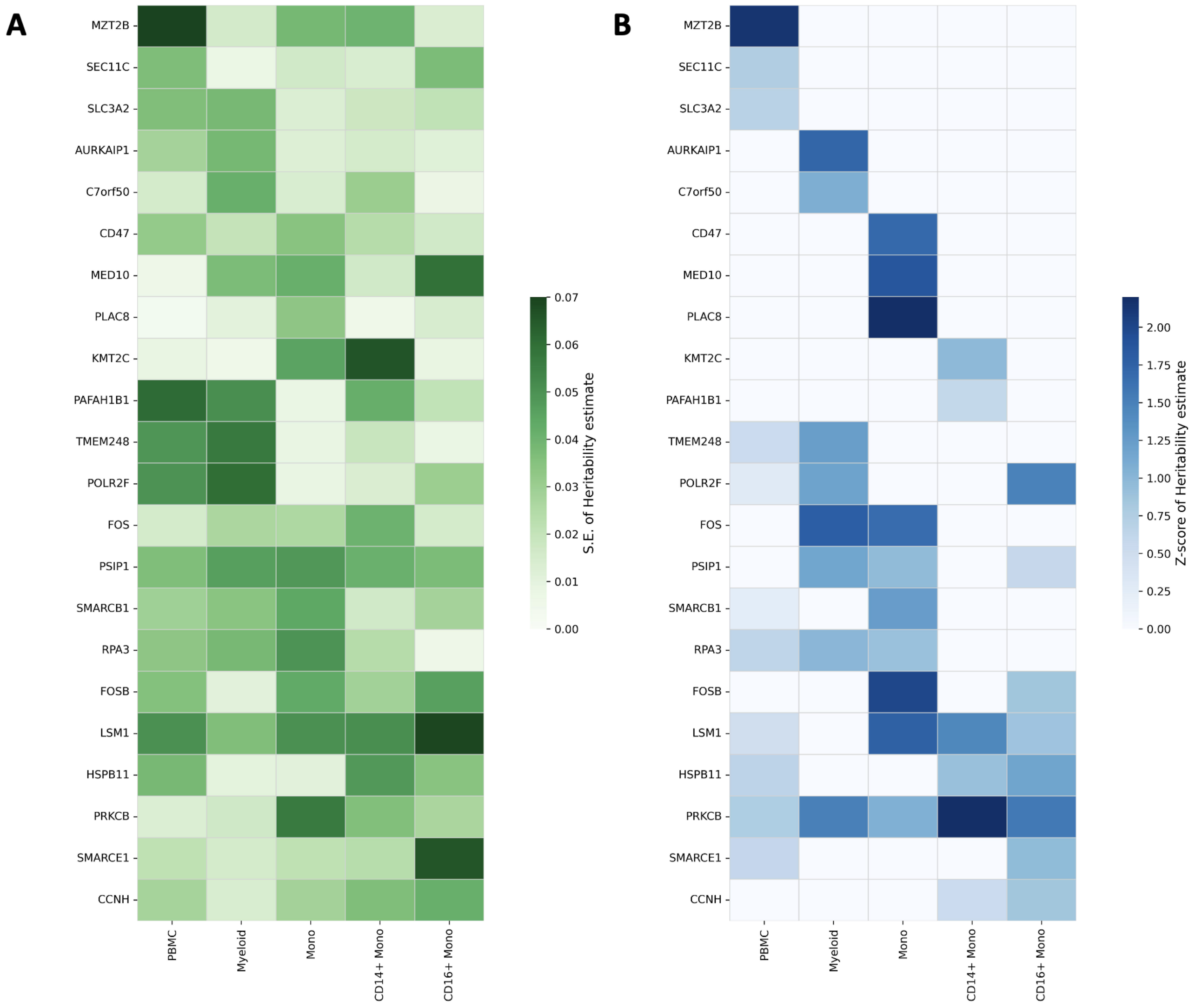


**Supplementary Figure 9: Standard error and z-score of *cis*-heritability estimates across varying granularities of immune cell subtypes.** Standard error of scGeneHE *cis*-heritability estimates for immune-relevant genes across the myeloid lineage corresponding to **Figure 5** (**a**). Corresponding z-scores of scGeneHE *cis*-heritability estimates (**b**).

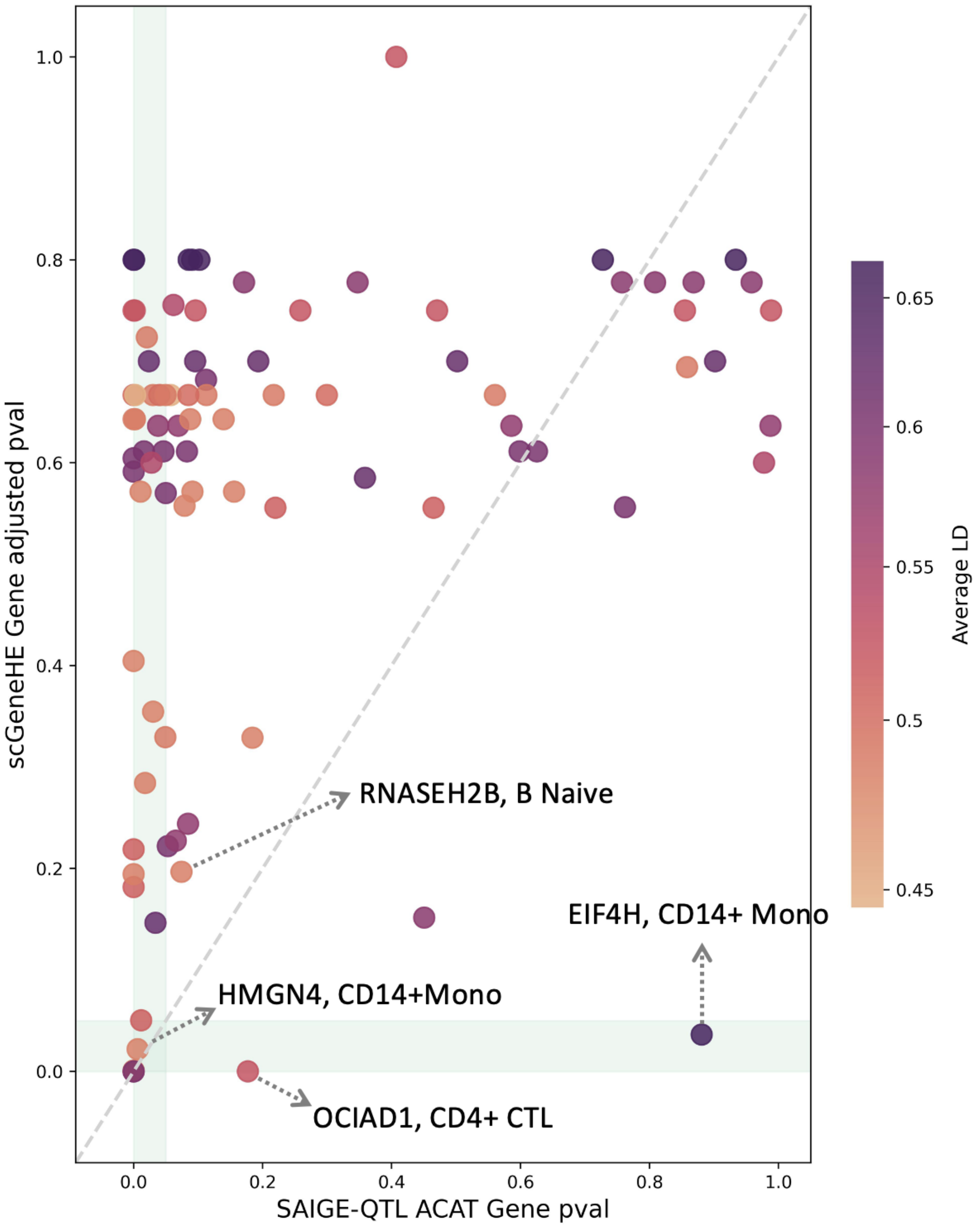


**Supplementary Figure 10: Comparison of gene-level statistics between scGeneHE and SAIGE-QTL ACAT-V test.** FDR-corrected p-values for *cis*-heritability from scGeneHE (y-axis) versus FDR-corrected gene-level p-values calculated by ACAT-V in SAIGE-QTL (x-axis). Vertical and horizontal light green bands indicate FDR $< 5\%$ for SAIGE-QTL and scGeneHE, respectively. Data is shown for genes exemplified in **Figure 4**. Color of dots indicates the average LD r2 within the *cis*-region of each gene.

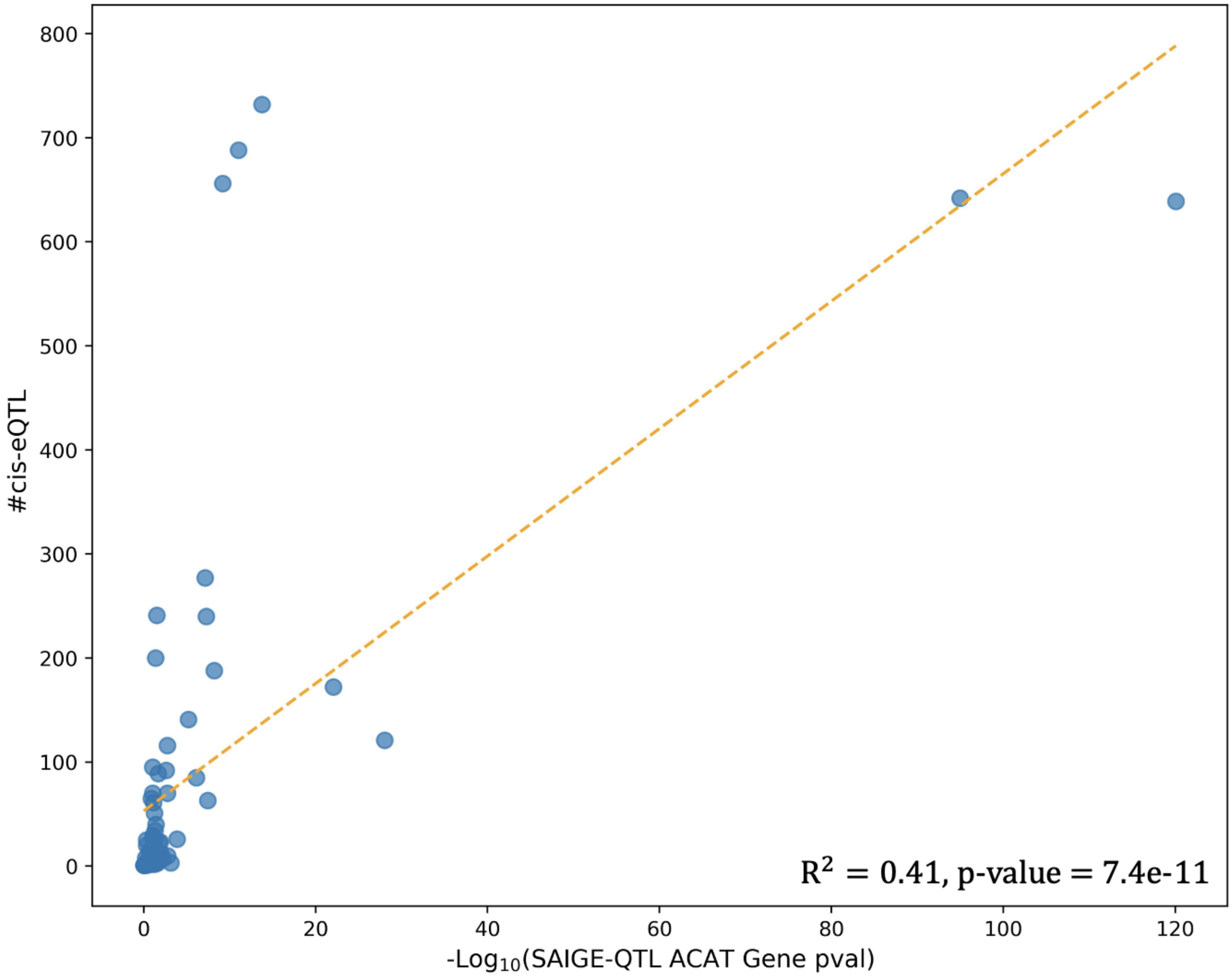


**Supplementary Figure 11: Comparison of ACAT-V gene-level p-values and number of *cis*-eQTLs for several pairs of genes and cell types.** Data corresponds to gene cell-type pairs from **Figure 6**. The dashed orange line is the line of best fit from ordinary least squares (OLS) regression. The squared correlation between -log10(ACAT gene p-value) and the number of *cis*-eQTLs from SAIGE-QTL is 0.41, suggesting that if LD can influence the number of significant cis-eQTLs, it likely affects ACAT-V test p-values as well.

### *Supplementary Table Legends*

**Supplementary Table 1: Numerical results for Figure 2A: point estimates of *cis*-heritability of scGeneHE and pseudobulk across varying sparsity rates and true heritability values in simulations.** Across 5 different true *cis*-heritability settings, we generate single cell and pseudobulk expression profiles for *cis*-heritable genes and corresponding null genes, with a donor sample size of 100, 5 causal eQTLs per gene, 50 cells per donor, and 80% sparsity rate. We evaluate *cis*-heritability estimation by applying scGeneHE to simulated single-cell expression and applying GCTA to corresponding pseudobulk profiles. We report the standard deviation across *cis*-heritability estimates for 100 independently simulated genes. 'lower' and 'upper' denotes the lower and upper bounds of the 95% confidence interval.

**Supplementary Table 2: Numerical results for Figure 2B: power and type I error rate of eGene detection for scGeneHE and pseudobulk across varying sparsity rates and true heritability values in simulations.** Across 5 different true *cis*-heritability settings, we evaluate the power of detecting truly *cis*-heritable genes and the type I error rate of incorrectly detecting null genes as *cis*-heritable. The following parameters are fixed: 100 donors, 50 cells per donor, 5 causal eQTLs per gene, and 80% sparsity rate. We report the standard deviation across power and type I error rates for 100 independently simulated genes.

**Supplementary Table 3: Numerical results for Figure 2C: sensitivity and specificity of eGene detection across varying true heritability values in simulations.** We generate 1000 uniformly distributed p-values and estimate the true positive (sensitivity) and false positive (1-specificity) rates at each p-value used to determine if a gene's *cis*-heritability is significantly non-zero. Columns with 'valid' as suffix represent the number of converged simulations.

**Supplementary Table 4: Numerical results for Figure 3B: *cis*-heritability estimates from scGeneHE and pseudobulk across different cellular heterogeneity levels and sparsity rates in simulations.** We evaluate *cis*-heritability estimates of *cis*-heritable genes and null genes with varying degrees of shared cellular environment and sparsity rates. The following parameters are fixed: 10% true heritability, 100 donors, 50 cells per donor, and 5 causal eQTLs per gene. We report the standard deviation across *cis*-heritability estimates for 100 independently simulated genes. 'lower' and 'upper' denotes the lower and upper bounds of the 95% confidence interval.

**Supplementary Table 5: Numerical results for Figure 3C: power of eGene detection by scGeneHE and pseudobulk across different cellular heterogeneity levels and sparsity rates in simulations.** Across 5 different true *cis*-heritability settings, we evaluate the power of detecting significant *cis*-heritable genes using scGeneHE and the pseudobulk approach. The following parameters are fixed: 10% true heritability, 100 donors, 50 cells per donor, and 5 causal eQTLs per gene. We report the standard deviation across power rates for 100 independently simulated genes.

**Supplementary Table 6: Meta-data on GWAS summary statistics implicating 166 genes mapped to loci associated with rheumatoid arthritis from the GWAS Catalog.** Meta-data includes study accessions, type of arthritis, ancestry of population cohort, significance of GWAS association, implicated loci, genotype sequencing platform, and other fields.

**Supplementary Table 7: Numerical results for Figure 4: *cis*-heritability estimates of 166 genes across 11 cell types in OneK1K cohort across scGeneHE, pseudobulk, and bulk approaches.** We generate *cis*-heritability estimates using scGeneHE and GCTA for pseudo-bulk and bulk from GTEx whole blood tissue[17,18]. We report the scGeneHE estimate of the *cis*-genetic component and cell environment component as $tau_1$ and $tau_2$, respectively. The standard error of scGeneHE is calculated from 100 bootstraps of data from each gene-cell-type pair. GCTA generates analytical standard errors for pseudobulk and bulk heritability estimates. FDR-correction is conducted per-gene across cell types.

**Supplementary Table 8: Numerical results for Figure 5: scGeneHE *cis*-heritability estimates of 200 genes across varying granularities of immune subtypes.** We generate *cis*-heritability estimates using scGeneHE across 200 randomly generated genes in PBMC, myeloid, monocyte, CD14+ monocyte, and CD16+ monocyte cell types. scGeneHE's estimate of the genetic component and cell environment component are reported by $tau_1$ and $tau_2$, respectively. Z-scores and standard errors of scGeneHE are calculated from 100 bootstraps of data from each gene-cell-type pair.

**Supplementary Table 9: Numerical results for Figure 6: gene-level p-value estimates using scGeneHE and SAIGE-QTL ACAT-V test and *cis*-eQTL association analysis results.** We generate gene-level p-values using the ACAT-V test as used by SAIGE-QTL and compare to scGeneHE *cis*-heritability p-values. FDR-corrected p-values are also reported for both methods. We calculate the number of significant eQTLs with correction across all gene-cell-type pairs. The *cis*-region and average linkage disequilibrium within the *cis*-region of each gene are also reported.
